# High-throughput differentiation of human blood vessel organoids reveals overlapping and distinct functions of the cerebral cavernous malformation proteins

**DOI:** 10.1101/2024.12.04.626588

**Authors:** Dariush Skowronek, Robin A. Pilz, Valeriia V. Saenko, Lara Mellinger, Debora Singer, Silvia Ribback, Anja Weise, Kevin Claaßen, Christian Büttner, Emily M. Brockmann, Christian A. Hübner, Thiha Aung, Silke Haerteis, Sander Bekeschus, Arif B. Ekici, Ute Felbor, Matthias Rath

**Affiliations:** Department of Human Genetics, University Medicine Greifswald and Interfaculty Institute of Genetics and Functional Genomics, University of Greifswald, Greifswald, Germany; Department of Dermatology and Venerology, Rostock University Medical Center, Rostock, Germany; ZIK plasmatis, Leibniz Institute for Plasma Science and Technology (INP), Greifswald, Germany; Institute of Pathology, University Medicine Greifswald, Greifswald, Germany; Institute of Human Genetics, Jena University Hospital, Friedrich Schiller University, Jena, Germany; Department of Human Medicine, MSH Medical School Hamburg, Hamburg, Germany; Institute of Human Genetics, Friedrich-Alexander-University (FAU) Erlangen-Nürnberg and Universitätsklinikum Erlangen, Erlangen, Germany; Institute for Molecular and Cellular Anatomy, University of Regensburg, Regensburg, Germany; Faculty of Applied Healthcare Science, Deggendorf Institute of Technology, Deggendorf, Germany; Institute for Molecular Medicine, MSH Medical School Hamburg, Hamburg, Germany; Organoid Expertise Center, MSH Medical School Hamburg, Hamburg, Germany

**Keywords:** blood vessel organoids, human induced pluripotent stem cells, CRISPR/Cas9 genome editing, cerebral cavernous malformations, single-cell RNA sequencing

## Abstract

Cerebral cavernous malformations (CCMs) are clusters of thin-walled enlarged blood vessels in the central nervous system that are prone to recurrent hemorrhage and can occur in both sporadic and familial forms. The familial form results from loss-of-function variants in the *CCM1*, *CCM2*, or *CCM3* gene. Despite a better understanding of CCM pathogenesis in recent years, it is still unclear why *CCM3* mutations often lead to a more aggressive phenotype than *CCM1* or *CCM2* variants. By combining high-throughput differentiation of blood vessel organoids from human induced pluripotent stem cells (hiPSCs) with a *CCM1*, *CCM2*, or *CCM3* knockout, single-cell RNA sequencing, and high-content imaging, we uncovered both shared and distinct functions of the CCM proteins. While there was a significant overlap of differentially expressed genes in fibroblasts across all three knockout conditions, inactivation of *CCM1*, *CCM2*, or *CCM3* also led to specific gene expression patterns in neuronal, mesenchymal, and endothelial cell populations, respectively. Taking advantage of the different fluorescent labels of the hiPSCs, we could also visualize the abnormal expansion of *CCM1* and *CCM3* knockout cells when differentiated together with wild-type cells into mosaic blood vessel organoids. In contrast, *CCM2* knockout cells showed even reduced proliferation. These observations may help to explain the less severe clinical course in individuals with a pathogenic variant in *CCM2* and to decode the molecular and cellular heterogeneity in CCM disease. Finally, the ability to differentiate blood vessel organoids in a 96-well format will further facilitate their use in drug discovery and other biomedical research studies.

**STATEMENTS AND DECLARATIONS:** *Conflicts of interest statement:* The authors declare no competing interests. The here described protocol for high-throughput organoid synthesis has been filed as a patent application at the European Patent Office (Process number: EP24213596.0)

*Author contribution statement:* MR, DSk, and UF designed the study. DSk, VS, LM, and RAP performed most of the functional experiments. SH and TA performed the CAM assays. SR performed the immunohistochemical stainings. SB, DSi, DSk, and VS performed the confocal microscopy and high-content imaging analyses. AE, CB, and EMB performed the scRNA sequencing analysis. AW and CAH performed and analyzed the karyotyping of the hiPSC clones. DSk, RAP, VS, KC, MR, and SB analyzed the data. DSk, VS, LM, and MR prepared figures. All authors contributed to the interpretation of the results. DSk, RAP, VS, and MR drafted the manuscript, and all authors contributed to writing.

*Ethics statement:* This study does not involve human participants or animal subjects.

*Availability of data and materials:* All relevant data are published within the paper and the supplementary files. ScRNA sequencing data can be accessed through the Gene Expression Omnibus (GEO) database (record number: GSE276497).

## INTRODUCTION

Cerebral cavernous malformations (CCMs), sometimes also called cavernomas, cavernous angiomas, or cavernous hemangiomas, are leaky vascular lesions in the brain and spinal cord that can cause seizures, intracranial hemorrhages (ICH), or non-hemorrhagic focal neurological deficits (NH-FND) [1,2]. With a prevalence of approximately 1 in 200, CCM is one of the most common cerebrovascular diseases. Although most CCMs are sporadic, there is also a familial form of CCM that accounts for 6 to 7 % of all cases [3]. Since the first description of a CCM family by H. Kufs in 1928 [4] and the identification of the three disease genes *CCM1* (also known as *KRIT1*), *CCM2*, and *CCM3* (also known as *PDCD10*) between 1999 and 2005 [5-9], CCM research has developed rapidly. Extensive studies in different cell culture models, human CCM tissue samples, and mouse, zebrafish, and other model organisms have contributed significantly to our current understanding of CCM pathogenesis [10-12]. It is now well established that although familial CCM is a classic autosomal dominant disorder, the inactivation of *CCM1*, *CCM2,* or *CCM3* is recessive at the cellular level [11]. The deregulation of key signaling pathways that initiate CCM formation, e.g., gain of MEKK3-KLF2/4 signaling or increase of RhoA/ROCK activity [13,14], is only triggered by the biallelic inactivation of a *CCM* gene. Furthermore, next-generation sequencing of bulk and single nucleus DNA from human CCMs has shown that the proportion of cells with a biallelic *CCM* gene variant is often rather low within the vascular lesions [15-17]. While *in vivo* and *in vitro* research has also led to the first phase 1 and 2 clinical trials, no pharmacological CCM therapy has yet been approved for clinical use [18-20]. Thus, neurosurgical treatment often remains the only option for symptomatic CCMs, especially after a previous hemorrhage or when they cause medically refractory seizures [1]. Since surgical resection is always associated with the risk of postoperative morbidity, the development of new CCM drugs remains a top priority in research. The increasing availability of advanced *in vitro* methods is raising hopes in many fields of biomedical sciences to significantly accelerate drug discovery. Organoid cultures, one of these advanced *in vitro* methods, have already enabled a wide range of new applications in basic and translational research, regenerative medicine, and drug discovery [21]. Using specific growth factors, small molecules, extracellular matrix components, and cell culture conditions, human pluripotent stem cells can be very effectively differentiated into brain, lung, kidney, liver, and other specific organoid types [22]. For example, the differentiation of human blood vessel organoids, first described by Wimmer and colleagues [23,24], now provides new insights into the complex processes of vasculogenesis and angiogenesis in health and disease. Especially in combination with CRISPR/Cas9 genome editing in fluorescently labeled human induced pluripotent stem cells (hiPSCs), human blood vessel organoids can help to better understand how wild-type (WT) and knockout (KO) cells interact within the CCM lesions. This aspect of CCM pathogenesis is still poorly understood but could represent a promising target to suppress CCM development or progression [25]. Using mosaic WT/KO blood vessel organoids differentiated from mEGFP-tagged *CCM3* KO and mTagRFPT-tagged *CCM3* WT hiPSCs, we recently directly visualized a significantly increased KO cell proliferation [26]. This abnormal *in vitro* behavior perfectly reflects the *in vivo* situation, where a clonal dominance of *Ccm3* KO endothelial cells was observed in CCM mouse models with a Confetti fluorescence reporter system [27,28]. Whether *CCM1* and *CCM2* KO cells also show such oncogenic properties is yet unknown.

Despite their many advantages, vascular organoid cultures have been challenging to scale up, e.g., for high-throughput studies. To address this limitation, we have established a protocol that eliminates the most labor-intensive step of manually extracting vascular networks from the extracellular matrix, allowing the differentiation of human blood vessel organoids in a 96-well format. Furthermore, the protocol does not require animal collagen I and bovine calf serum. Using this high-throughput compatible and nearly xeno-free protocol in combination with scRNA sequence analysis, we demonstrate that blood vessel organoids are powerful tools for CCM research, providing deep insight into the cellular and molecular consequences of *CCM1*, *CCM2,* or *CCM3* inactivation.

## MATERIALS AND METHODS

### Cell culture

Human induced pluripotent stem cells (hiPSCs) were cultured in 6-well plates (Greiner Bio-One, Frickenhausen, Germany, #657160) using Essential 8 Flex medium (Thermo Fisher Scientific, Waltham, MA, USA, #A2858501). Cell culture dishes were pre-coated with growth factor-reduced Matrigel (Corning Inc., Corning, NY, USA, #356231). HiPSCs were thawed and plated at a density of 10,000 cells/cm^2^ in Matrigel-coated 6-well plates in the presence of RevitaCell supplement (1:100) (Thermo Fisher Scientific, #A2644501). HiPSC cultures were regularly tested negative for mycoplasma contamination by PCR. The hiPSC lines AICS-0036-006 [WTC-mEGFP-Safe harbor locus (AAVS1)-cl6 (mono-allelic tag); hPSCreg ID: UCSFi001-A-12; RRID:CVCL_JM19], AICS-0054-091 [WTC-mTagRFPT-CAAX-Safe harbor locus (AAVS1)-cl91 (mono-allelic tag); hPSCreg ID: UCSFi001-A-23; RRID:CVCL_VK84] and AICS-0016-184 [WTC-mEGFP-ACTB-cl184 (mono-allelic tag); hPSCreg ID: UCSFi001-A-3; RRID:CVCL_JM16], which are part of the Allen Cell Collection (Allen Institute for Cell Science, Seattle, WA, USA), were purchased from the Coriell Institute (Camden, NJ, USA). *CCM1* and *CCM2* knockout (KO) AICS-0036-006 hiPSCs as well as *CCM1, CCM2,* and *CCM3* KO AICS-0016-184 hiPSCs were generated in this study following our established protocols [26,29]. The generation of *CCM3* KO AICS-0036-006 hiPSCs has been described earlier [26]. "*CCM1* KO", "*CCM2* KO", and "*CCM3* KO" hereafter refer to KO cells derived from AICS-0036-006-hiPSCs unless otherwise specified. For CRISPR/Cas9 genome editing, single guide RNA (sgRNA): Cas9 ribonucleoprotein complexes with the following target sequences were used: *CCM1* 5′-GGAGCTCCTAGACCAAAGTA-3′; *CCM2* 5′-GGTCAGTTAACGTCCATACC-3′; *CCM3* 5′-CAACTCACCTCATTAAACAC-3′ (Integrated DNA Technologies, Coralville, IA, USA). The introduction of homozygous or compound heterozygous loss-of-function variants by CRISPR/Cas9-editing in the generated clones was verified by amplicon-based next-generation sequencing. HiPSC quality control was performed by karyotyping and immunofluorescence analysis of the pluripotency markers OCT4, SSEA4, SOX2, and TRA-1-60 using the Pluripotent Stem Cell 4-Marker Immunocytochemistry Kit (Thermo Fisher Scientific, #A24881) as previously described [29].

### Generation of hiPSC, mesodermal, and vascular aggregates

For the generation of hiPSC aggregates and their differentiation into vascular aggregates, the previously described protocol of Wimmer and colleagues [23,24] was modified to allow aggregate formation in a 96-well format. Briefly, on day -2, hiPSCs were detached and dissociated using StemPro Accutase (Thermo Fisher Scientific, #A1110501). Cells were counted and diluted to a concentration of 1,000 cells/well in fresh aggregation medium [23,24] containing 50 μM Y-27632 (Stemcell Technologies, Vancouver, Canada, #72304). Cells were seeded in 100 μL medium per well in ultra-low attachment PrimeSurface 96 Slit-well plates (Sbio, Hudson, NH, USA, #MS9096SZ). Plates were centrifuged at 300 x g for 3 min at room temperature and incubated at 37 °C and 5 % CO_2_ for 48 hours to promote effective hiPSC aggregation. On day 0, the PrimeSurface 96 Slit-well plates were washed by adding 25 mL of N2B27 medium to the plate using a 25 mL serological pipette. The plate was gently tilted to all sides to distribute the medium evenly. The medium was removed by titling the 96-slit-well plate and slowly aspirating it from one corner of the plate with a 25 mL serological pipette. Next, 25 mL of N2B27 medium + 12 µM CHIR99021 (Tocris Bioscience, Bristol, United Kingdom, #4423) + 30 ng/mL BMP-4 (Miltenyi Biotec, Bergisch Gladbach, Germany, #130-111-164) were added. The cells were incubated at 37 °C and 5 % CO_2_ for 72 hours to induce mesodermal differentiation. On day 3, 25 mL of the old medium were aspirated, the aggregates were washed as described above, and 25 mL of N2B27 medium + 100 ng/mL VEGF-A (Peprotech, Hamburg, Germany, #100-20-50) + 2 µM forskolin (Miltenyi Biotec, #130-117-341) were added. The cells were incubated at 37 °C and 5 % CO_2_ for 48 hours to induce vascular differentiation. To study aggregation efficiency KO and WT hiPSCs were seeded with different cell densities ranging from 200 cells/well to 1800 cells/well. Following the protocol of mesodermal and endothelial differentiation described above, the aggregates were imaged on days 0, 3, and 5 (= hiPSC, mesodermal, and vascular aggregates, respectively). The imaging was performed using an EVOS M5000 imaging systems microscope (Thermo Fisher Scientific). The size of the aggregates was determined using FIJI v.1.54 [30]. The aggregates were fixed with 4 % PFA on day 5. The following primary and secondary antibodies were used for immunofluorescent evaluation: ZO-1 (Novus Biologicals, Centennial, CO, USA, 1:100, #NBP1-85047), anti-rabbit IgG Alexa 647 (Thermo Fisher Scientific, 1:200 #A21246).

### Generation of human blood vessel organoids

For the generation of vascular networks and blood vessel organoids, the previously described protocol by Wimmer and colleagues [23,24] was modified as follows: On day 5, vascular aggregates were transferred to a 96-well Akura plate (InSphero, Schlieren, Switzerland, #CS0900403) and embedded in a human collagen I/Matrigel matrix. The following amounts of human collagen I (Advanced BioMatrix, Carlsbad, CA, USA, #5007-20ML) and Matrigel solution were used for one 96-well Akura plate: 490 µL each for the first and second matrix layers (367,5 µL of human collagen I solution + 122.5 µL of Matrigel). Matrigel and human collagen I were kept on ice. To form the first matrix layer, 3 µL of human collagen I/Matrigel solution was added to each well of an empty Akura 96-well plate. The plate was incubated at 37 °C for 2 h to solidify the matrix, while a second Akura plate was prepared as a transfer plate. To avoid air bubbles at the bottom of the transfer plate, the wells were washed with StemPro-34 serum-free medium (StemPro-34 SFM, Thermo Fisher Scientific, #10639011). After removing 10 mL of the N2B27 medium from the PrimeSurface 96 Slit-well plate containing the vascular aggregates, a multichannel pipette with cut pipette tips was used to aspirate the vascular aggregates in 70 µL medium from the bottom of the Slit-Well plate. They were transferred to the transfer plate, and excess medium was removed. 50 µL of StemPro-34 SFM was then added to each well before 40 µL of medium containing the aggregates was transferred to the Akura plate containing the first collagen I/Matrigel matrix layer using a multichannel pipette with cut pipette tips. Once again, as much medium as possible was aspirated before 5 µL of human collagen I/Matrigel solution was slowly added as a second matrix layer embedding the aggregates. For solidification of the matrix, the plate was incubated for 2 hours at 37 °C. Meanwhile, StemPro-34 SFM containing 15 % Panexin CD (PAN Biotech, Aidenbach, Germany, #P04-930500) was prepared in a Falcon tube and heated to 37 °C in a water bath for 30 minutes. 80 µL of StemPro-34 SFM supplemented with 15 % Panexin CD, 100 ng/mL FGF-2 (Miltenyi Biotec, #130-093-564), and 100 ng/mL VEGF-A were carefully added to the center of each well. The plate was incubated at 37 °C and 5 % CO_2_ to induce sprouting. The medium was replaced on days 7 and 10. On day 12, the matrix plugs with the vascular networks were transferred without manual extraction from the gel to individual wells of a PrimeSurface 96 Slit-well plate by pipetting. Medium was changed as previously described by adding 25 mL of StemPro-34 SFM supplemented with 15 % Panexin CD, 100 ng/mL FGF-2, and 100 ng/mL VEGF-A to the Slit-well plate. Over the next two days, the networks were regularly pipetted up and down with cut pipette tips to promote the dissolution of excess matrix. The medium was exchanged again on day 14. On day 17, the blood vessel organoids were ready for following analyses. To generate mosaic blood vessel organoids, mTagRFPT-tagged AICS-0054 WT hiPSCs and *CCM1* KO, *CCM2* KO, or *CCM3* KO mEGFP-tagged AICS-0036 hiPSCS were mixed in a 19:1 ratio and differentiated to blood vessel organoids as described above. For immunofluorescence analysis, vascular networks and blood vessel organoids were fixed in 1 % PFA for 30 min or 1 h, respectively. Antibody staining was performed as described elsewhere [23]. The following primary and secondary antibodies were used: CD31 (RnD Systems, Minneapolis, MN, USA, #BBA7, 1:50), PDGFR-β (Cell Signaling Technology, Danvers, MA, USA, #3169S, 1:50), VE-cadherin (Cell Signaling Technology, #2500S, 1:400), and collagen IV (Novus Biologicals, #NB120-6586, 1:100), anti-Mouse IgG Fluor 350 (Thermo Fisher Scientific, #A-11045), anti-Rabbit IgG Alexa Fluor 647 (Thermo Fisher Scientific, 1:200, #A21246), anti-Mouse IgG Alexa 555 (Abcam, Cambridge, England, 1:200, #ab150114). Nuclei were stained with Hoechst 33342. Imaging was performed using an Operetta CLS imaging system (PerkinElmer, Waltham, MA, USA) in non-confocal mode or a Stellaris 8 (Leica, Wetzlar, Germany) microscope.

### Perfusion of human blood vessel organoids in OrganoPlates Graft plates

*In vitro* perfusion assays were performed with OrganoPlate Graft plates (Mimetas, Oegstgeest, Netherlands, #6401-400-B). Gel preparation was performed according to the manufacturer’s instructions. 3 µL of gel containing 4 mg/mL rat tail Collagen I (Ibidi, Gräfelfing, Germany, #50201), 100 mM HEPES (Thermo Fisher Scientific, #15630106), and 3.7 mg/mL NaHCO_3_ (Merck, Darmstadt, Germany, #6329) were added to the gel inlet of the OrganoPlate. For polymerization of the matrix, the plate was incubated for 15 minutes at 37 °C and 5 % CO_2_. Human umbilical vein endothelial cells (HUVECs, PromoCell, Heidelberg, Germany) were cultured in endothelial cell growth medium (ECGM, PromoCell) supplemented with 10 % fetal bovine serum (FBS) (Thermo Fisher Scientific, #A5670701). Following the manufacturer’s instructions, 2 μL of 1[×[10^4^ cells/μL were seeded into the perfusion inlets of the Organoplate. After adding 50 µL of medium to the perfusion inlets, the plate was incubated for 2-3 hours at 37 °C and 5 % CO_2_. After cell attachment, 50 µL of medium was added to the perfusion outlets, and the plate was placed on an interval MIMETAS rocker with an inclination of 14° and an interval of 8 minutes. Medium was replaced every two to three days. Cells were cultured for three days until tube formation was completed. To initiate sprouting, 50 µL of medium mixed with an angiogenic cocktail of 37.5 ng/mL VEGF-A, 37.5 ng/mL FGF-2, 37.5 ng/mL rhMCP-1 (ImmunoTools, Friesoythe, Germany, #11343384), 37.5 ng/mL rhHGF (ImmunoTools, #11343413), 250 nM S1P (Sigma Aldrich, St. Louis, MO, USA, #S9666) and 37.5 ng/mL PMA (Sigma Aldrich, # P1585) were added to the graft chamber according to the manufacturer’s instruction. To graft hiPSC-derived vascular networks in the OrganoPlates, medium was removed from all channels after six days of sprouting. Next, perfusion inlets and outlets were filled with ECGM supplemented with 10 % FBS. StemPro-34 SFM containing 15 % PCD and the previously described angiogenic cocktail was added to the graft chamber. Finally, the vascular networks were transferred into the graft chamber using cut pipette tips. Medium was replaced every two to three days. After five days, the perfusion of the vascular network was visualized by adding medium containing 0.5 mg/mL 150 kDA tetramethylrhodamine (TMR)-amino dextran (Fina Biosolutions LLC, Rockville, MD, USA, #TMR-Amdex150K) to the left perfusion inlet and outlet. After 15 minutes, the plate was fixed using 4 % PFA for 30 minutes and imaged according to the manufacturer’s instructions.

### Grafting of organoids on chicken chorioallantoic membranes (CAM model)

To study perfusion of the orgnaoids in the chorioallantoic membrane (CAM) 3D in vivo model, complete blood vessel organoids were resuspended in StemPro-34 SFM medium containing 15 % PCD and 10 % DMSO, frozen in liquid nitrogen, and shipped overnight. Fertilized chicken eggs were rinsed with water, brushed, and disinfected with 70 % ethanol. Afterward, they were immediately placed on a rotation device inside a ProCon egg incubator (Grumbach, Asslar, Germany) at a constant temperature of 37.8 °C, a pCO_2_ of 5 %, and with the humidity calibrated to 63 %. On the fourth day of incubation, two windows were cut into the eggshell [31-33]. The first window, that was located at the air entrapment of the egg, enabled an equilibrium of pressure inside the egg and was permanently closed with Leukosilk (BSN Medical, Hamburg, Germany). The second window was cut into the longitudinal part of the egg and covered with a removable strip of Leukosilk. After organoid engraftment, the eggs were placed on glass trays containing autoclaved sand in the non-rotating compartment of the egg incubator. After a one-week growth period of the organoids on the CAM, they were explanted and prepared for histological analysis. First, the organoids were immersion fixed in 4 % paraformaldehyde (PFA) under constant movement for 7 days at 4 °C. After removing the PFA, the organoids were washed in PBS (phosphate buffered saline, pH 7,4) for 30 min under constant movement. This procedure was repeated three times. Consequently, the samples were dehydrated and embedded in paraffine. Tissue samples of a thickness of 6 µm were cut using a microtome and stained with hematoxylin and eosin (H&E staining).

### Single-cell RNA sequencing (scRNA-Seq)

After blood vessel organoid synthesis, ten organoids per genotype were pooled, and enzymatic dissociation was performed. First, blood vessel organoids were transferred to a sterile Petri dish and minced with a scalpel. Second, the minced organoids were then suspended in 1 mL enzymatic mix containing 4 mg Liberase TH solution (Sigma Aldrich), 30 mg Dispase II (Sigma Aldrich), and 0.5 mg DNase I solution (New England Biolabs, Ipswich MA, USA) in PBS and incubated at 37 °C and 5 % CO_2_ for 30 minutes. Third, the dissociated blood vessel organoids were mixed and passed through a 70 µm cell strainer. 3 mL of DMEM/F-12 (Thermo Fisher Scientific, #11330032) with 10 % FBS (Thermo Fisher Scientific, #A5670701) was added to stop the enzymatic reaction. Next, cells were centrifuged at 300 × g for 5 minutes, resuspended in StemPro-34 medium with 10 % DMSO, and frozen in liquid nitrogen for shipment. Library preparation for scRNA-Seq was performed with the Chromium Single Cell Next GEM 3’ Reagent Kit v3.1 (10x Genomics, Pleasanton, CA, USA) on a Single Cell 3’ device following manufacturer’s instructions. Sequencing was performed on a Novaseq 6000 sequencer (Illumina, San Diego, CA, USA). Cellranger version 7.0.1 was used to generate read data, demultiplex single cells, perform sequence alignment, and generate raw count data. Further filtering of raw data, generation of clusters, and analysis of gene expression differences were performed with Seurat 4.0.4. In brief, cells with a UMI count under 2500 and less than 600 detected genes were removed, and the mitochondrial DNA and ribosomal RNA cut-off values were set to 10 % and 5 %, respectively. Seurat SCTransform was further used to normalize samples. Data was integrated using a mutual nearest neighbors (MNN) approach and unsupervised cell clustering was performed with the Leiden algorithm. Web-based Cell-type-Specific Enrichment Analysis of Genes (WebCSEA) was applied to identify enriched cell types of generated clusters [34] using the protein-coding marker genes with a log_2_FC ≥ 0.5 for each cluster. For each cluster, The CZ CELLxGENE Discover browser [35] was used to visualize the tissue and cell type-specific expression levels of the top 10 marker genes in specific clusters. Differentially expressed genes (DEG) of individual clusters were analyzed with the ShinyGO tool [36] and subjected to gene set enrichment analyses with the GO cellular process and molecular function gene sets.

### Differentiation of hiPSCs into ECs and endothelial co-culture

*CCM1* KO AICS-0016 hiPSCs, *CCM3* KO AICS-0016 hiPSCs, WT AICS-0016 hiPSCs, and WT AICS-0054 hiPSCs were differentiated into ECs (iECs) using the STEMdiff Endothelial Differentiation Kit (Stemcell Technologies) as previously described [29]. To confirm endothelial differentiation, cells were fixed with 4 % PFA at passage 1 and stained for the endothelial markers CD31 (Cell Signaling 3528S, 1:800), VE-cadherin (2500S, 1:400, Cell Signaling) and VWF (Thermo Fisher Scientific, MA5-14029, 1:66). For endothelial co-culture assays, AICS-0016 KO iECs were mixed with AICS-0054 WT iECs in a 1:9 ratio and cultured together for 6 days in EndoGRO-MV media or STEMdiff Endothelial Expansion media on 96-well plates. Co-cultures of AICS-0016 WT iECs and AICS-0054 WT iECs were used as controls. Wells were fixed with 4 % PFA, and nuclei were stained with Hoechst 33342 (Thermo Fisher Scientific, #62249). FIJI was used to compare the area of mEGFP-tagged cells with the total area.

### Statistical analysis

Statistical analysis was performed in GraphPad Prism 8 (GraphPad Software, USA). Normality was assessed using the Shapiro-Wilk test. For normally distributed data, statistical significance was evaluated using multiple two-sample t-tests with Welch’s correction to account for unequal variances between groups. For non-normally distributed data, pairwise Mann-Whitney U-tests were performed, as described in the figure legends. Holm-Šídák adjustment for multiple testing was applied within each outcome category, with the following notation: ns = P ≥ 0.05; * = P < 0.05; ** = P < 0.01; *** = P < 0.001.

## RESULTS

### High-throughput-compatible and nearly xeno-free synthesis of human blood vessel organoids

To reduce the workload of blood vessel organoid differentiation and make it more suitable for high-throughput applications, we first modified some key steps (Fig. 1A,B) of the original differentiation protocol [23,24]. The use of Slit-well plates allowed precise control of the aggregate size while reducing the time spent on media exchange compared to conventional ultra-low attachment 6-well or 96-well cell culture plates. Since all wells in a Slit-well plate share one media pool, medium exchange can be performed using a 25 mL serological pipette (Fig. 1C). Our new protocol also eliminates the most labor-intensive step of blood vessel organoid differentiation, namely cutting out the vascular networks from the collagen I/matrigel matrix under a sterile bench. Since the embedding of vascular aggregates on day 5 of the protocol takes place in special Akura 96 spheroid microplates, which have a conical well design with a bottom diameter of only 1 mm, the vascular networks can be directly transferred by pipetting to a new 96-well plate after sprouting. The amount of excess matrix around the vascular networks is minimal (Fig. 1B,C) and easily dissolved by regular pipetting up and down with cut pipette tips over the next seven days. The use of these special cell culture plates makes it relatively easy to learn the new protocol (Fig 1D). Besides modifying the workflow, we also adapted the reagents used in the differentiation and replaced bovine collagen I and fetal bovine serum (FBS) with human collagen I and human platelet lysate (hPL) or Panexin CD (PCD), respectively. Since FBS, hPL, and PCD led to comparable sprouting efficiencies (Fig. 1E), we decided to use the chemically defined serum replacement PCD for further experiments. This means that the differentiation protocol is now nearly xeno-free. Only Matrigel could not be replaced. Vascular networks and blood vessel organoids generated with the modified protocol are approximately 0.5 to 1.0 mm in size and display highly complex endothelial networks with associated pericytes (Fig. 1F,G). Using the OrganoPlate Graft technology, we were able to demonstrate proper lumen formation in hiPSC-derived vascular networks. After grafting mEGFP-tagged vascular networks onto a HUVEC bed, perfusion was visualized by introducing TMR-amino-dextran (Fig. 1H).

**Fig. 1.**
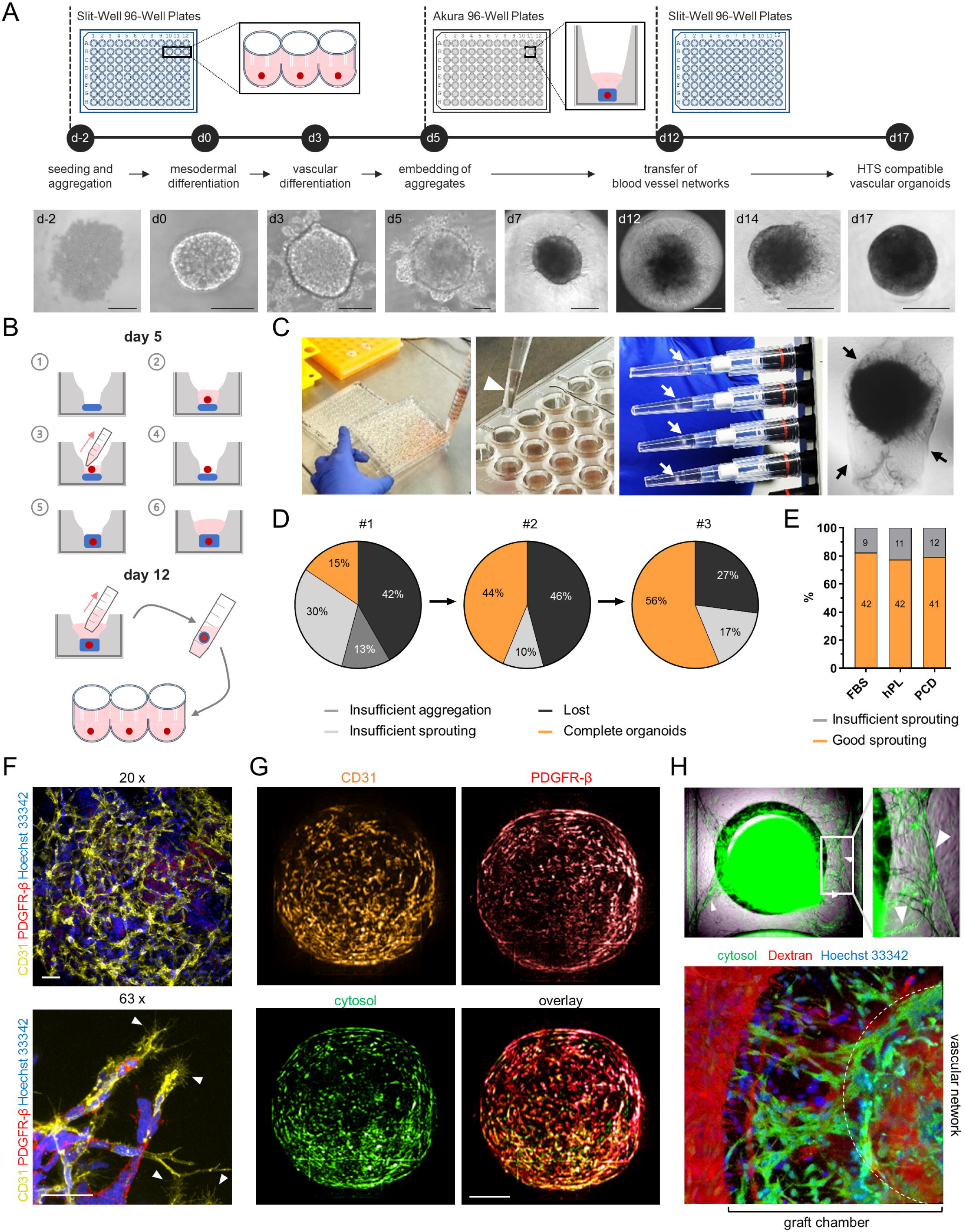
High-throughput (HT)-compatible and nearly xeno-free synthesis of vascular networks and blood vessel organoids from fluorescently tagged human induced pluripotent stem cells (hiPSCs). **A** Schematic illustration of the new differentiation protocol and representative images for the main differentiation steps (scale bars: d-2, d0, d3, d5 = 100 µm; d8, d12 = 250 µm; d14, d17 = 500µm). **B** Shown are the steps of embedding the vascular aggregates in an Akura 96-well plate and transferring the vascular networks from the Akura 96-well plate to a PrimeSurface 96 Slit-well plate. **C** The use of PrimeSurface 96 Slit-well plates reduces the time required for medium exchange (left image). Akura 96-well plates allow aggregates to be embedded in small cavities, minimizing the matrix surrounding the vascular networks (black arrows) and allowing direct transfer of vascular networks (white arrows) to new plates without time-consuming manual extraction of the networks from the gel (middle and right images). **D** The new protocol is simple to handle and achieves high synthesis efficiency after minimal training. Shown are the efficiencies of three training runs. **E** The sprouting efficiency is maintained when fetal bovine serum (FBS) is replaced with human platelet lysate (hPL) or chemically defined Panexin CD (PCD). The total numbers of sufficiently sprouted networks and vascular aggregates with insufficient sprouting are written inside the bars. **F,G** HiPSC-derived vascular networks (F) and blood vessel organoids (G) differentiated with the HT-compatible and nearly xeno-free protocol consist of a complex network of endothelial cells (CD31) and associated pericytes (PDGFR-β) [representative images; scale bars: 50 µm (F); 200 µm (G)]. White arrowheads indicate angiogenic sprouts. **G** Perfusion of vascular networks with TMR-amino-dextran in OrganoPlate graft plates shows anastomoses between the GFP-labeled vascular networks and the HUVEC-derived vascular bed (top, white arrowheads) as well as correct formation and permeability of the vascular networks (bottom).

### Perfusion of human blood vessel organoids on chorioallantoic membranes (CAM)

As a next step, we wanted to combine our easy to use, scalable, and almost xeno-free blood vessel organoid differentiation protocol with the chorioallantoic membrane (CAM) 3D in vivo model, which is commonly used for perfusion approaches [33,37]. One week after grafting onto the CAM, the blood vessel organoids were explanted for hematoxylin and eosin (H&E) staining and immunohistochemical staining for CD31 and PDGFR-β (Fig. 2A-C). We found intact endothelial and pericyte structures in the grafted blood vessel organoid sections. To differentiate between chicken and human blood vessels, an antibody that specifically targets human CD31 (PECAM1) was used. In H&E-stained tissue sections, nucleated chicken erythrocytes were found within the vascular structures of the blood vessel organoids, indicating successful perfusion in the CAM model (Fig. 2D).

**Fig. 2.**
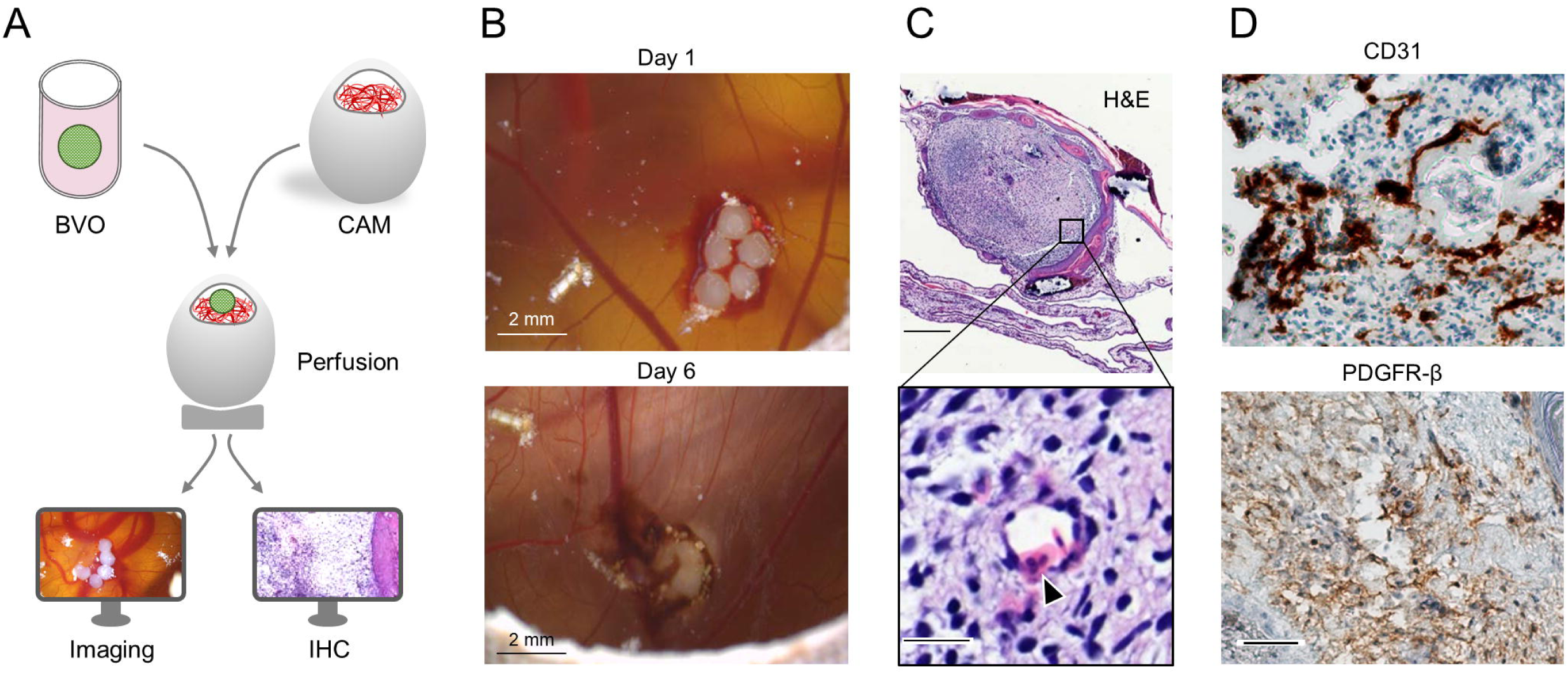
Perfusion of blood vessel organoids (BVO) on chorioallantoic membranes (CAM). **A** Schematic illustration of the perfusion approach. **B** Shown are blood vessel organoids cultivated on the CAM associated with chicken blood vessels. The pictures were taken on day 1 and day 6 (scale bars = 2 mm). **C** Sectioning and H&E staining demonstrated nucleated chicken erythrocytes within the vascular structures of the blood vessel organoid (upper scale bar = 500 µm; bottom scale bar = 25 µm). **D** The expression of CD31 (upper image) and PDGFR-β (lower image) were verified by immunohistochemistry staining (brown) (scale bar = 100 µm).

### Irregular aggregation of hiPSCs, mesodermal, and vascular endothelial cells upon *CCM1* and *CCM3* inactivation

To learn more about the pathobiology of CCM, we generated *CCM1* and *CCM2* KO hiPSCs from the mEGFP-tagged AICS-0036-006 hiPSC line using CRISPR/Cas9 genome editing and limiting dilution cloning (Fig. S1). The generation of mEGFP-tagged *CCM3* KO AICS-0036-006 hiPSCs had already been described in our previous publication [26]. Comparing KO and WT cells in the first steps of differentiation into blood vessel organoids already revealed some differences (Fig. 3A,B). *CCM1* KO and *CCM3* KO aggregates were significantly larger than the wild-type controls. These differences were observed at various cell seeding densities and became more pronounced over time at the mesodermal (day 3) and vascular differentiation (day 5) stages (Fig. 3B,C,E). In contrast, no significant size differences were found for *CCM2* KO aggregates (Fig. 3B,D). Hypothesizing that impaired cell-cell-contacts in *CCM* KO vascular aggregates might be an explanation for this observation, we stained for the tight junction protein 1 (= zonula occludens protein 1/ZO-1) often described as impaired in CCM literature [38-41]. Imaging revealed that *CCM1* and *CCM*3 KO vascular aggregates contained large cell-free cavities that were not or only barely present in WT or *CCM2* vascular aggregates, respectively (Fig. 3F). This is a reasonable explanation for the differences in size between the genotypes.

**Fig. 3.**
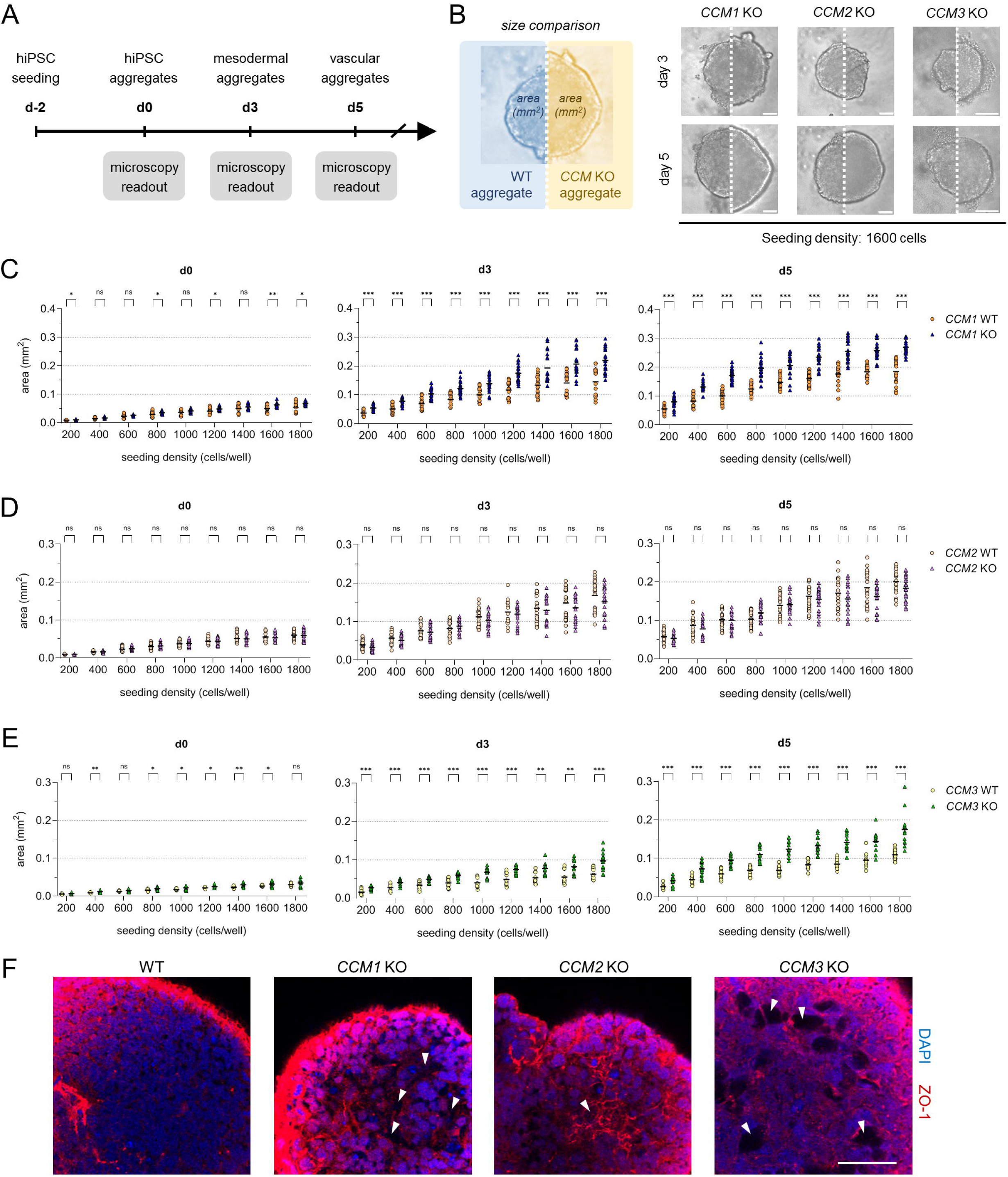
Size differences between WT and KO aggregates. **A** Schematic illustration of the initial hiPSC aggregation and the induction of mesodermal and vascular differentiation. **B** Shown are representative images of wild-type (WT; left half of each image) and *CCM1*, *CCM2,* and *CCM3* knockout aggregates (KO; right half of each image) on days 3 and 5 (initial seeding density: 1600 cells/well, scale bars: 100 µm). **C-E** WT and *CCM1* (C), WT and *CCM2* (D), WT and *CCM3* (E) KO hiPSCs were seeded with variable seeding densities (200 to 1800 cells/well) at d-2. The cross-sectional area of the aggregates was determined on days 0, 3, and 5. Shown are individual data points, line represents mean, n=12-24 per genotype in three independent biological replicates. Multiple two-sample t-tests with Welch’s correction and Holm-Šídák adjustment for multiple testing were used for statistical analyses (ns = Padj ≥ 0.05; * = Padj < 0.05; ** = Padj < 0.01; *** = Padj < 0.001). **F** Staining of tight junction protein 1 (= zonula occludens protein 1/ZO-1) revealed significant structural differences between WT and *CCM1*, *CCM2,* and *CCM3* KO vascular aggregates (Scale bars: 50 µm). White arrows indicate cell free cavities.

### Single-cell RNA sequencing (scRNA-seq) reveals different cellular compositions of WT and KO blood vessel organoids

To further characterize the cellular composition and genotype-specific gene expression differences in WT and KO blood vessel organoids, we used scRNA-seq analysis (Fig. 4A). After integrating the datasets and unsupervised clustering, we identified 11 clusters with unique gene expression signatures. The visualization in an UMAP plot demonstrated good separation of the clusters (Fig. 4B). By analyzing cell-type-specificity of marker genes using the WebCSEA tool, we identified clusters of ECs (c1, c9), perivascular/stromal cells (c3, c5, c6, c10), cells with neuronal and glial gene expression signature (c7, c8, c11), and proliferating cells (c4) (Fig. 4B,C and S3). Perivascular/stromal cell clusters were characterized by fibroblastic, mesenchymal, and smooth muscle cell gene expression signatures (Fig. S3). Inactivation of *CCM1*, *CCM2,* or *CCM3* resulted in significant shifts in the cellular composition of the organoids. While there were many overlapping effects, comparative analysis also revealed genotype-specific differences (Fig. 4D-F). For example, *CCM1* KO organoids had a marked over-representation of cells in clusters c3, c9, and c10. In contrast, *CCM2* KO organoids showed a remarkably high proportion of cells in cluster c5. Substantial changes in cellular composition were also observed in *CCM3* KO organoids. However, we did not find specific over- or underrepresented clusters that distinguished *CCM3* from *CCM1* or *CCM2* KO organoids.

**Fig. 4.**
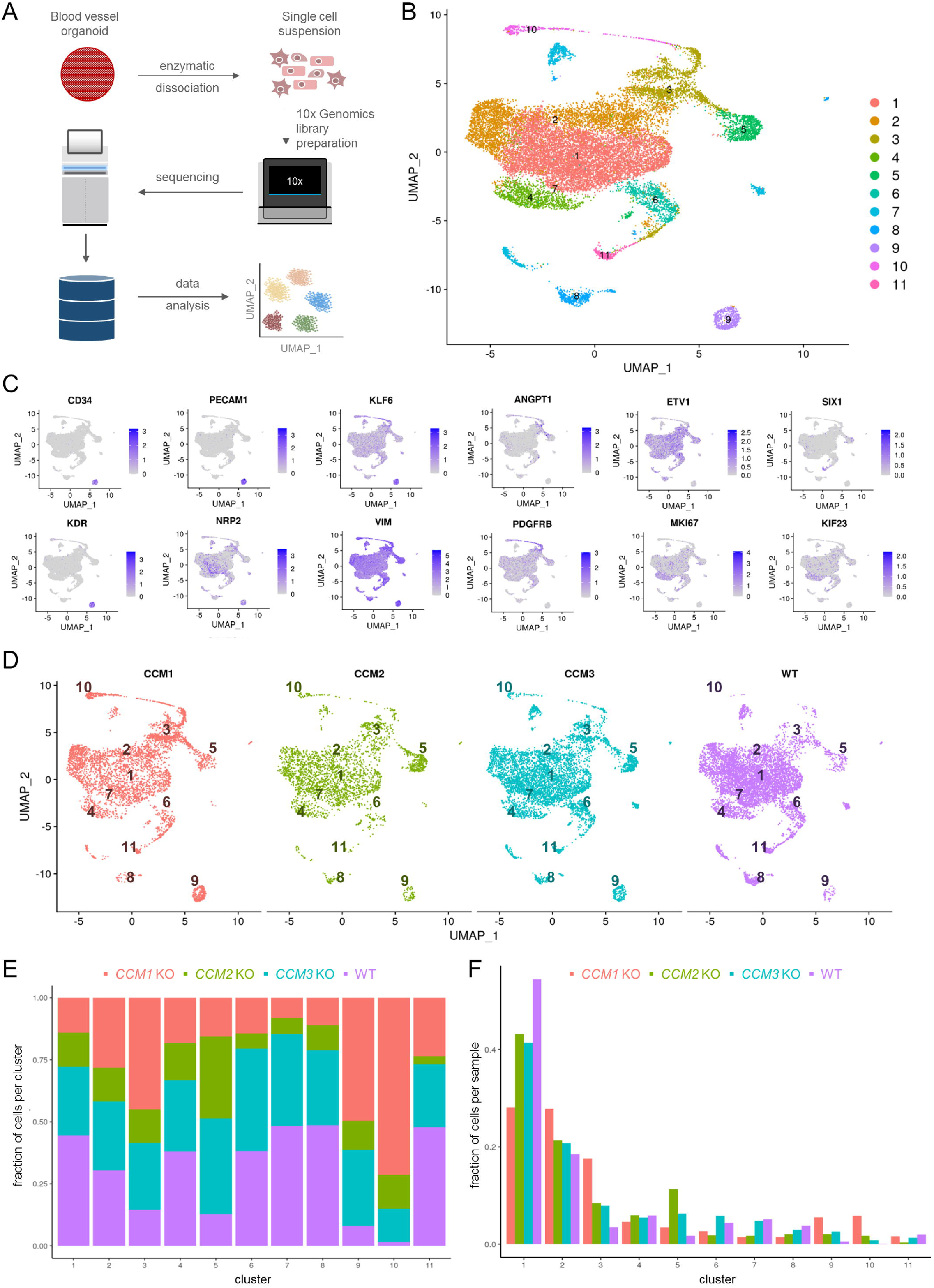
Changes in cellular composition of KO blood vessel organoids. **A** Experimental design of the single cell RNA sequencing (scRNA-seq) experiments for KO and WT organoids. **B** Uniform Manifold Approximation and Projection (UMAP) visualization of the scRNA-seq data with cells colored according to the unsupervised clustering results. **C** UMAP plots colored by the expression levels of marker genes predominantly expressed in endothelial, pericyte, smooth muscle and proliferating cells. Purple color indicates high expression. Low expression is indicated by grey color. **D** Separate UMAP plots of scRNA-seq data from *CCM1* KO, *CCM2* KO, *CCM3* KO, and WT blood vessel organoids. The numbers represent the different clusters. **E** The composition of the clusters is illustrated by the proportion of *CCM1* KO, *CCM2* KO, *CCM3* KO, and WT cells within each cluster. **F** The cellular composition of KO and WT blood vessel organoids is illustrated by the distribution of *CCM1* KO, *CCM2* KO, *CCM3* KO, and WT samples into the different clusters.

Overlapping and genotype-specific effects were also observed at the gene expression level (Fig. 5A). Clusters c3, c5, c6, c8, c9, and c10 showed the highest numbers of differentially expressed genes (DEGs) per genotype (Fig. 5B-D). Indeed, cluster c10 had the highest number of DEGs (Fig. 5B-D). Since this cluster also showed the largest overlap of DEGs between the three KO conditions (Fig 5E,F), it was classified as the main overlap cluster (Fig. 5G). Among the most up- or downregulated genes in this fibroblast cell cluster (Fig. 6A and S3) were *EBF1*, *COL3A1,* and *DLK1*, which were similarly deregulated in all three KO genotypes (Fig. 6B-D). A GO ontology analysis of the 51 genes significantly upregulated in all three KO conditions showed a strong enrichment for genes that are involved in collagen fibril organization (Fig. 6E). A pathway analysis for the 55 DEGs downregulated in all three KO conditions revealed a more diffuse picture with weaker enrichment for different signaling pathways (Fig. 6F).

**Fig. 5.**
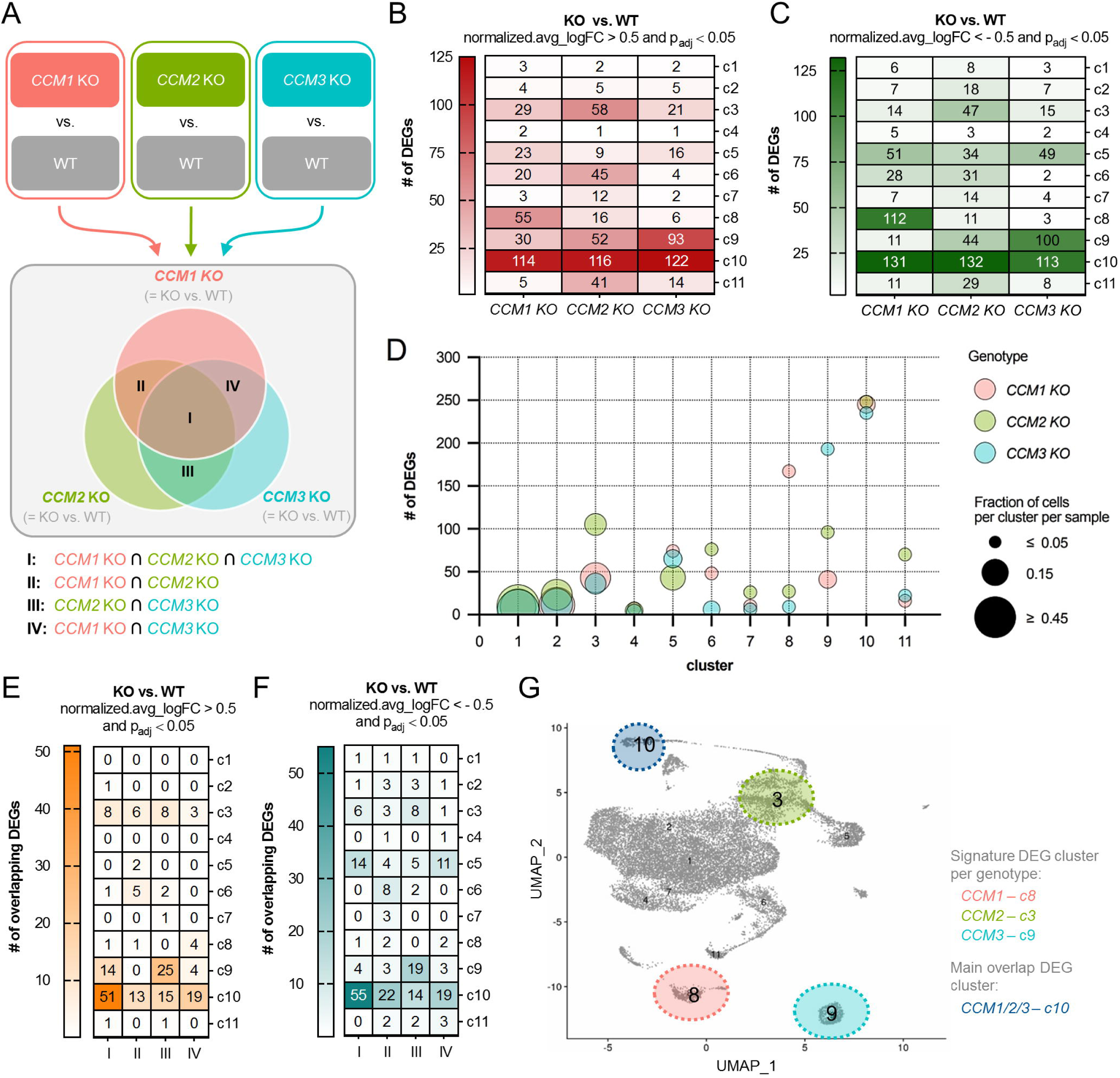
Overlapping and genotype-specific gene expression differences in knockout blood vessel organoids. **A** Scheme of the analysis strategy and the selected comparisons. **B,C** Heatmaps with the numbers of significantly up- (B) and downregulated (C) genes per cluster and genotype. DEGs **=** differentially expressed genes. **D** The relationship between the total number of DEGs per genotype and cluster and the fraction of cells per cluster in the *CCM1*, *CCM2,* and *CCM3* KO samples is illustrated in a bubble plot. **E,F** Heatmaps with the numbers of overlapping up- (E) and downregulated (F) DEGs between the different genotypes. I = *CCM1* KO ∩ *CCM2* KO ∩ *CCM3* KO; II = *CCM1* KO ∩ *CCM2* KO; III = *CCM2* KO ∩ *CCM3* KO; IV = *CCM1* KO ∩ *CCM3* KO. **G** The main overlap DEG cluster (c10) and the signature DEG clusters per genotype (*CCM1* KO: c8; *CCM2* KO: c3; *CCM3* KO: c9) are highlighted in the UMAP plot. Significantly up- and downregulated genes were defined as those with a normalized.avg_logFC (KO vs. WT) > 0.5 and p_adj_ < 0.05 or with a normalized.avg_logFC (KO vs. WT) < -0.5 and p_adj_ < 0.05, respectively.

**Fig. 6.**
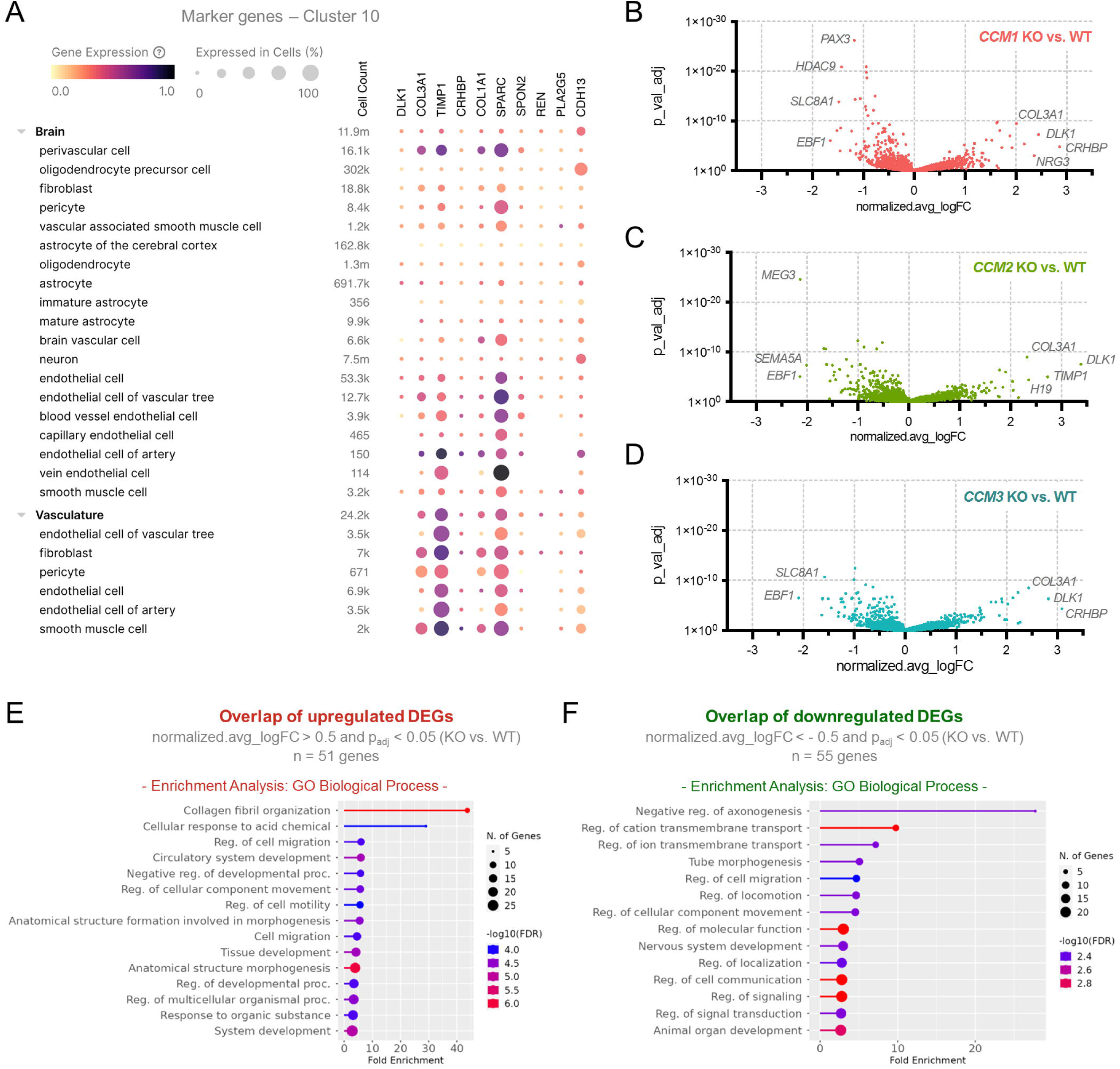
Overlapping gene expression differences in cluster 10. **A** The CZ CELLxGENE Discover browser was used to visualize the tissue and cell type-specific expression levels of the top 10 marker genes identified in cluster 10. Shown are cell types that are typically found in the brain and the vasculature. Purple color indicates high expression. Low expression is indicated by yellow color. The percentage of cells of the specific cell type that express the marker gene is visualized by the size of the circles. **B**-**D** Genotype-specific gene expression differences in cluster 10 are shown in Volcano plots for *CCM1* (B), *CCM2* (C), and *CCM3* (D) KO samples. **E,F** Overlapping up- (E) and downregulated (F) genes were subjected to gene set enrichment analyses with the GO biological process gene set. Significantly up- and downregulated genes were defined as those with a normalized.avg_logFC (KO vs. WT) > 0.5 and p_adj_ < 0.05 or with a normalized.avg_logFC (KO vs. WT) < -0.5 and p_adj_ < 0.05, respectively. Overlapping DEGs were defined as DEGs found in *CCM1*, *CCM2,* and *CCM3* KO samples (= *CCM1* KO ∩ *CCM2* KO ∩ *CCM3* KO).

In the other clusters, the numbers of overlapping DEGs were significantly lower. We even found genotype-specific signatures in certain clusters. For example, only *CCM1* KO organoids had a high number of DEGs in cluster c8. Similar characteristic DEG clusters were also found in *CCM2* and *CCM3* KO organoids. Here, specific gene expression differences were found in clusters c3 and c9, respectively (Fig 5E-G). In the EC cluster c9, a total of 193 DEGs were found in the *CCM3* KO organoids, but only 41 and 96 in *CCM1* and *CCM2* KO organoids, respectively. Only 18 DEGs were found to overlap in all three KO genotypes (Fig. S4A,B). These were mainly associated with angiogenesis-related processes (Fig. S4C). There was additional overlap between *CCM2* and *CCM3* KO organoids but hardly any between *CCM1* and *CCM2* or *CCM1* and *CCM3* KO organoids (Fig. 5E,F). The *CCM3*-specific DEGs in this cluster (Fig. S4D,E) mainly included genes involved in cell-matrix adhesion (Fig. S4F,G). The *CCM1*-specific DEGs in the neuronal-like cell cluster c8 were associated with axonogenesis and neuron differentiation, and the *CCM2*-specific DEGs in the mesenchymal-like cell cluster c3 were associated with mesenchyme development and regulation of cell motility (Fig. S5 and S6).

### Abnormal proliferation of *CCM1*, *CCM2*, and *CCM3* KO cells in mosaic blood vessel organoids and endothelial co-cultures

We have previously demonstrated that *CCM3* KO cells show an abnormally high proliferation in WT/*CCM3* mosaic blood vessel organoids [26]. With our new differentiation protocol, we not only reproduced our previous result but also studied the behavior of *CCM1* KO and *CCM2* KO cells in mosaic blood vessel organoids. Wild-type hiPSCs labeled with mTagRFPT were mixed with mEGFP-labeled knockout hiPSCs (19:1 ratio) and differentiated into mosaic blood vessel organoids. In WT/*CCM3* KO and WT/*CCM1* KO mosaic blood vessel organoids, significantly increased mEGFP signals were observed (Fig. 7A,B). In WT/*CCM2* KO mosaic organoids, on the other hand, we saw significantly lower mEGFP signals (Fig. 7A,B). To better understand the interaction of KO and WT cells, we synthesized mosaic vascular networks and performed immunostaining analyses. A striking finding was that *CCM3* KO mosaic vascular networks appeared to be composed of much more convoluted CD31 positive endothelial and PDGFR-β positive pericyte networks than *CCM1* and *CCM2* KO mosaic networks (Fig. S7A). Another interesting finding was that VE-cadherin-positive endothelial cells appeared less abundant in mEGFP-labelled *CCM* KO cells, particularly *CCM3* KO cells (Fig. S7B and Supplementary Videos 1-4).

**Fig. 7.**
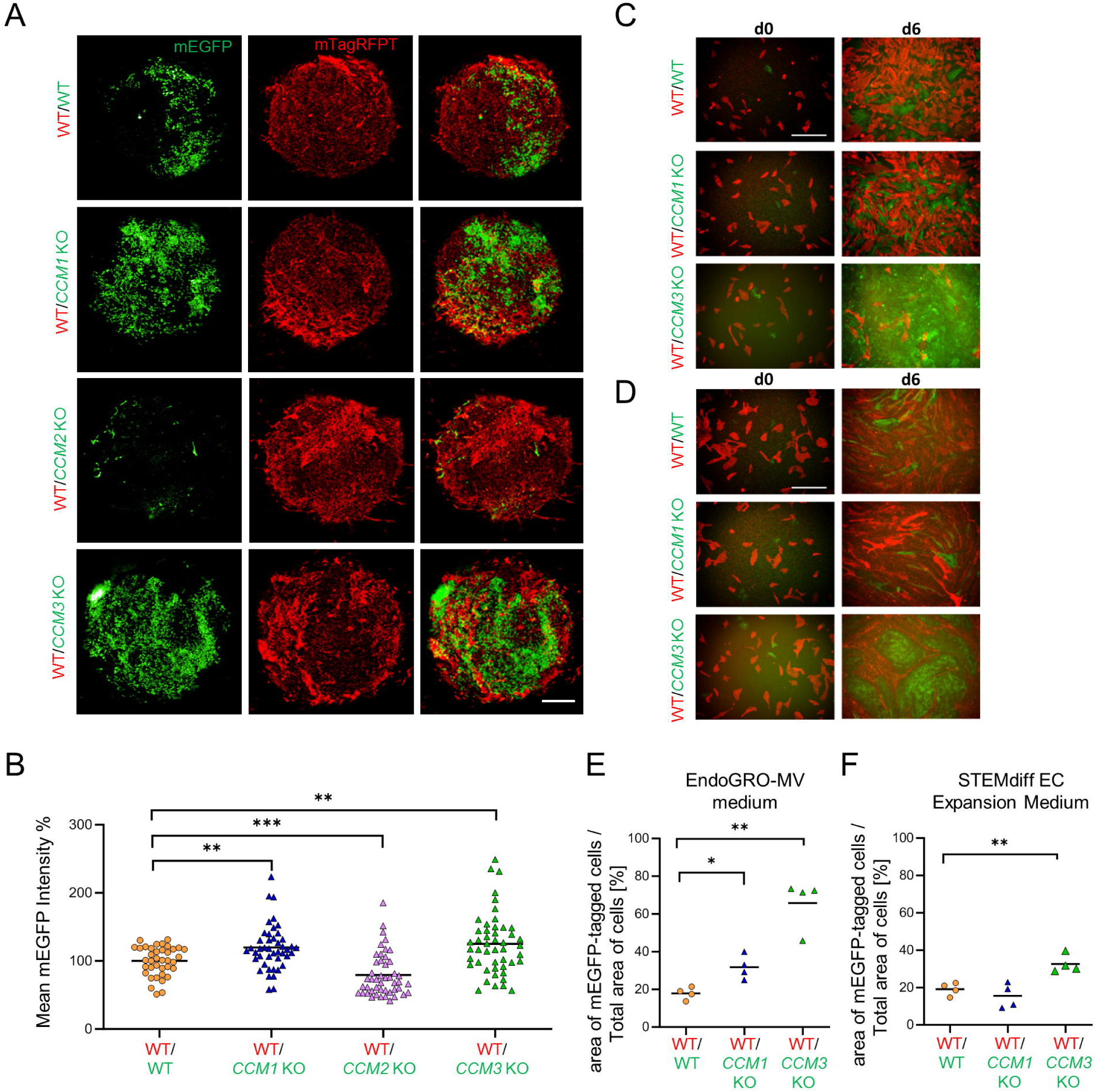
Abnormal proliferation of knockout cells in mosaic blood vessel organoids and EC co-cultures. **A** In mosaic WT/*CCM3* KO and WT/*CCM1* KO blood vessel organoids, an abnormally increased proliferation of *CCM1* KO and *CCM3* KO cells was observed. In mosaic WT/*CCM2* KO blood vessel organoids, a reduced proliferation of *CCM2* KO cells was found. KO cells were labeled with mEGFP. WT cells were labeled either with mTagRFPT or mEGFP (scale bar: 200 µm). **B** Mean mEGFP intensities in WT/KO mosaic organoids were normalized to the mean mEGFP intensity in WT/WT mosaic organoids. Mosaic blood vessel organoids were differentiated in three independent runs (n= 38-48 per genotype). Data are presented as individual data points and means. Statistical significance was assessed using the Mann-Whitney U-test with Welch’s correction (**P < 0.01, ***P < 0.001). **C,D** In a 2D co-culture validation approach, mEGFP-labeled *CCM1* or *CCM3* KO iECs and mTagRFPT-labeled WT iECs were seeded in a 1:9 ratio and cultivated in either EndoGRO-MV (C) or STEMdiff EC expansion medium (D) (scale bar: 200 µm). **E,F** After 6 days, the area of mEGFP-labeled cells compared to all cells was determined using FIJI software (n = 4 per genotype in three independent biological replicates). Data are presented as individual data points and means. Multiple two-sample t-tests with Welch’s correction and Holm-Šídák adjustment for multiple testing were used for statistical analyses (* = Padj < 0.05; ** = Padj < 0.01).

To confirm the observed proliferative advantage of *CCM3* KO and *CCM1* KO cells in mosaic blood vessel organoids we went back to a two*-*dimensional iEC co-culture assay and mixed mTagRFPT-labeled WT iECs with mEGFP-labeled *CCM1* or *CCM3* KO iECs in a 9:1 ratio, which also showed significantly increased mEGFP signals after 6 days. Especially for *CCM3* KO iECs, a highly increased and almost "tumor-like" proliferation was observed in co-culture. This behavior was most striking in EndoGRO-MV medium (Fig. 7C,E). In STEMdiff endothelial expansion medium, the proliferation of *CCM3* KO iECs was also significantly increased, but the effect was less pronounced (Fig. 7D,F). Compared to *CCM3* KO iECs, *CCM1* KO iECs co-cultured with WT iECs in EndoGRO-MV medium showed a less pronounced proliferation advantage (Fig. 7C,E). In STEMdiff medium, no proliferation difference was found for *CCM1* KO iECs (Fig. 7D,F). Taken together, these data show that inactivation of *CCM1* or *CCM3* alone does not induce hyperproliferation but can trigger an abnormal proliferation in a specific microenvironment.

## DISCUSSION

Although the differentiation of blood vessel organoids from hiPSCs is still a relatively new cell culture technique [23], it already had a significant impact on endothelial and vascular biology research in recent years [24,42-44]. However, one limitation of the protocol has been its lack of scalability. In this study, we have simplified the workflow and present a high-throughput compatible protocol fully optimized for a 96-well format. Together with the use of a chemically defined, animal-free serum replacement and the possibility to perfuse the organoids or vascular networks on a CAM or in an OrganoPlate Graft plate, the new protocol facilitates the replacement of animal studies in biomedical research with organoid cultures. The latter two perfusion models can complement another recently described *in vitro* perfusion approach using a sophisticated microfluidic device [45] so that both simple and more complex perfusion experiments can now be realized. Since the vascularization and perfusion of organoids is also a high priority in other fields of biomedical research [46-48], these results may have an impact beyond vascular medicine.

Our data also demonstrate that blood vessel organoids are excellent tools for studying CCM disease. While mouse models have provided invaluable insights into key mechanisms of CCM pathogenesis, e.g., deregulation of MEKK3-KLF2/4 signaling, increased endothelial activation of PI3K-mTOR signaling, loss of junctional integrity in CCMs, clonal expansion of KO ECs, or recruitment of WT cells into growing CCM lesions [13,27,28,49-52], they can hardly be used for a comprehensive comparison between *CCM1*, *CCM2,* and *CCM3* KO conditions. In accordance with the 3R principle (Replacement, Reduction, Refinement) [53], the differentiation of blood vessel organoids from hiPSCs now allows the direct comparison of the cellular and molecular consequences of *CCM1*, *CCM2,* and *CCM3* gene inactivation in a complex three-dimensional cell culture system. Differences between the genotypes were already evident at the stages of mesodermal and vascular aggregates. In particular, *CCM1* and *CCM3* KO aggregates were significantly larger than WT and *CCM2* KO aggregates. Further evaluation revealed large cavities in *CCM1* and *CCM3* KO vascular aggregates, while *CCM2* KOs showed cavities to a lesser extent, providing an explanation for the size differences. These genotype-dependent morphological abnormalities observed in vascular aggregates are reminiscent of CCM lesions in *Ccm^BECKO^* mice. Yang and colleagues showed that overall lesion volume was the highest in *Ccm3*-deficient mice, followed by *Ccm1*-deficient mice, with *Ccm2*- deficient mice showing the least extensive lesion growth [54].

However, the differences between the three KO genotypes were not limited to the vascular aggregate stage of organoid synthesis. Consistent with data from two *Ccm3* mouse models [27,28], we have recently demonstrated tumor-like proliferation of *CCM3* KO cells in mosaic WT/KO blood vessel organoids [26]. Here, we have reproduced this effect with our new protocol and demonstrated a similar behavior for *CCM1* KO cells. Previously, there was only limited evidence from *CCM1* KO blood outgrowth endothelial cells (BOECs) that inactivation of the *CCM1* gene can promote the development of oncogenic properties in ECs [55]. While a previous observation that *CCM1* silencing impairs EC proliferation [56] seems to contradict the results of our mosaic organoid experiments, our data also show that the proliferation advantage of *CCM1* and *CCM3* KO cells is not an inherent effect of the gene inactivation but can be induced in a specific microenvironment. Indeed, different media significantly modulated the tumor-like proliferation of KO iECs in WT/KO co-cultures. *CCM1* KO iECs proliferated abnormally in EndoGRO MV medium, but not in STEMdiff medium. A less pronounced but still significant increase in cell growth was also observed for *CCM3* KO iEC in STEMdiff medium. These results highlight the critical role of growth factors and other external factors in developing endothelial dysfunction in *CCM1*- and *CCM3*-deficient cells, which may represent a therapeutic target.

Interestingly, *CCM2* KO cells showed a different proliferation behavior in mosaic WT/KO blood vessel organoids than *CCM1* and *CCM3* KO cells and even exhibited impaired cell growth. Consistent with this observation, previous studies have shown that silencing *CCM2* does not stimulate endothelial proliferation or can even induce a growth arrest in ECs [57,56]. Contrary to CCM1 and CCM3, CCM2 has a paralog, CCM2L, which can at least partially compensate for its loss. *In vivo*, CCM2L cannot completely block CCM formation in mice with endothelial-specific *Ccm2* deletion (*Ccm2^iECKO^*) but prevents a more severe phenotype that would result from a double knockout of *Ccm2* and *Ccm2l* [58]. At the molecular level, this compensatory effect is mediated by a partial blockade of MEKK3 signaling by CCM2L [58,59]. Given that MEKK3-KLF2/4 signaling is important for regulating endothelial proliferation and survival [50,60,61], inhibition of this pathway by CCM2L could explain the different behavior of *CCM2* KO cells.

Further insights into the distinct and overlapping functions of CCM proteins were also gained from our blood vessel organoid scRNA-seq data. As expected, deregulation of angiogenesis-related processes was a common feature found in ECs of all three KO genotypes. One of the consistently upregulated genes in EC cluster 9 was *KLF2*, highlighting the pivotal role of MEKK3-KLF2/4 signaling in CCM disease [13,62]. Furthermore, downregulation of the endothelial tip cell marker gene *ESM1* and upregulation of the endothelial-to-mesenchymal transition (EndMT) marker genes *S100A4* and *COL3A1* provide further evidence for the impact of EndMT on CCM pathogenesis [51,63,64]. However, an advantage of the CCM blood vessel organoid model is that it also incorporates the role of non-ECs. Indeed, in all three KO conditions, we identified many overlapping DEGs in the fibroblast cell cluster, which showed a strong enrichment of genes related to collagen fibril organization. Increased expression of extracellular matrix (ECM) genes in *CCM3* KO human brain microvascular pericytes and augmented deposition of fibronectin and collagen in mice with mural cell-specific deletion of *Ccm3* (*Ccm3*^smKO^) have previously been shown to contribute to the dissociation of ECs and pericytes in CCM lesions [65]. However, there also seems to be a cell non-autonomous regulation of ECM production by pericytes in CCM disease. For example, a study in mice with endothelial-specific deletion of *Ccm1* (*Ccm1^ECKO^*) revealed transforming growth factor-beta (TGFβ)-mediated upregulation of ECM genes in pericytes and increased fibronectin deposition in CCM lesions. [66]. The use of blood vessel organoids also revealed non-overlapping effects of *CCM1*, *CCM2*, and *CCM3* inactivation, which may explain differences in the severity of CCM disease in patients with pathogenic variants in these genes. While the aforementioned lack of tumor-like proliferation of *CCM2* KO cells may explain the milder course in patients with a *CCM2* mutation [67], the high number of *CCM3*-specific DEGs in EC cluster 9 indicates a more profound endothelial dysfunction, consistent with a more severe clinical phenotype in patients with a *CCM3* mutation [68,69]. Finally, the high number of *CCM1*-specific DEGs in a cell cluster with a neuronal-like gene expression signature gives a first idea of why seizures are the most common manifestation in patients with a *CCM1* mutation [67]. Although CCM1 is expressed in human neurons [70], future studies will have to show whether the gene expression differences are caused by cell-autonomous or non-autonomous mechanisms.

In summary, blood vessel organoids differentiated according to our novel high-throughput compatible protocol provide new insights into the complex CCM disease. Our results suggest that CCM research should focus not only on endothelial dysfunction, but also on the role of perivascular cells and tumor-like mechanisms to find a new CCM therapy.

## ACKNOWLEDGMENTS

This research was made possible through the Allen Cell Collection, available from Coriell Institute for Medical Research, which provided the hiPSC-lines. We would like to thank Nikolai Siemens, who is part of the Greifswald Imaging Facility, for assistance with the confocal microscopy.

## FUNDING

This work was funded by the German Federal Ministry of Education and Research (BMBF) [grant number 161L0276] and by the Research Network Molecular Medicine of the University Medicine Greifswald [FOVB-2020-01 and FOVB-2021-07]. MR was supported by a scholarship from the Gerhard Domagk program of the University Medicine Greifswald. High-content imaging analysis was supported by funding from the German Federal Ministry of Education and Research (BMBF, grant numbers 03Z22DN11 and 03Z22Di1, both to SB). DSi is financially supported by the Rostock Academy of Sciences (RAS).

## FIGURE LEGENDS

**Fig. S1** Quality control of unpublished CRISPR/Cas9 edited AICS-0016 and AICS-0036 hiPSC lines. *CCM1* KO, *CCM2* KO, and *CCM3* KO hiPSCs express the pluripotency markers TRA-1-60, OCT4, SOX2, and SSEA4 (representative images, scale bar: 200 µm). Karyotyping of knockout clones confirmed a normal karyotype (46, XY).

**Fig. S2** *CCM1* KO, *CCM2* KO, and *CCM3* KO iECs show regular expression of endothelial markers and reorganization of the actin cytoskeleton compared to wild-type controls. **A** Immunofluorescence analysis of the endothelial markers CD31, VE-cadherin, and VWF in WT and KO AICS-0016-derived iECs (scale bar = 75 µm). **B** Endogenously tagged actin in AICS-0016 lines reveals actin stress fiber formation in *CCM* KO conditions (scale bar = 25 µm).

**Fig. S3** Cell-type specificity analyses for scRNA-seq cell clusters. Results from WebCSEA showing the top 20 enriched general cell types for each cluster, which is summarized in the upper left panel. The results are based on the protein-coding marker genes with a normalized log_2_FC ≥ 0.5 for each cluster.

**Fig. S4** Gene expression differences in the *CCM3* signature cluster 9. **A** The CZ CELLxGENE Discover browser was used to visualize the tissue and cell type-specific expression levels of the top 10 marker genes identified in cluster 9. Shown are cell types that are typically found in the brain and the vasculature. Purple color indicates high expression. Low expression is indicated by yellow color. The percentage of cells of the specific cell type that express the marker gene is visualized by the size of the circles. **B,C** Overlapping DEGs (B) in cluster 9 were subjected to a gene set enrichment analysis with the GO biological process gene set (C). **D,E** Heatmaps of gene expression differences for significantly up- (D) and downregulated (E) genes found in *CCM3* KO, but not *CCM1* KO or *CCM2* KO samples (= *CCM3* specific DEGs). Shown are the normalized average logFC values (KO vs. WT) for the three genotypes. × = Genes without expression information in *CCM1* and *CCM2* KO samples. **F,G** *CCM3*-specific DEGs were subjected to gene set enrichment analyses with the GO cellular components (F) and molecular function (G) gene sets. Significantly up- and downregulated genes were defined as those with a normalized.avg_logFC (KO vs. WT) > 0.5 and p_adj_ < 0.05 or with a normalized.avg_logFC (KO vs. WT) < -0.5 and p_adj_ < 0.05, respectively.

**Fig. S5** Gene expression differences in the *CCM1* signature cluster 8. **A** The CZ CELLxGENE Discover browser was used to visualize the tissue and cell type-specific expression levels of the top 10 marker genes identified in cluster 8. Shown are cell types that are typically found in the brain and the vasculature. Purple color indicates high expression. Low expression is indicated by yellow color. The percentage of cells of the specific cell type that express the marker gene is visualized by the size of the circles. **B**-**D** Genotype-specific gene expression differences in cluster 8 are shown in volcano plots for *CCM1* (B), *CCM2* (C), and *CCM3* (D) KO samples. **E,F** The overlaps of upregulated (E) and downregulated (F) genes in *CCM1*, *CCM2*, and *CCM3* KO cells are shown as Venn diagrams. **G** *CCM1*-specific DEGs were subjected to a gene set enrichment analysis with the GO biological process gene set. Significantly up- and downregulated genes were defined as those with a normalized.avg_logFC (KO vs. WT) > 0.5 and p_adj_ < 0.05 or with a normalized.avg_logFC (KO vs. WT) < -0.5 and p_adj_ < 0.05, respectively.

**Fig. S6** Gene expression differences in the *CCM2* signature cluster 3. **A** The CZ CELLxGENE Discover browser was used to visualize the tissue and cell type-specific expression levels of the top 10 marker genes identified in cluster 3. Shown are cell types that are typically found in the brain and the vasculature. Purple color indicates high expression. Low expression is indicated by yellow color. The percentage of cells of the specific cell type that express the marker gene is visualized by the size of the circles. **B,C** The overlaps of upregulated (B) and downregulated (C) genes in *CCM1*, *CCM2*, and *CCM3* KO cells are shown as Venn diagrams. **F** *CCM2*-specific DEGs were subjected to a gene set enrichment analysis with the GO biological process gene set. Significantly up- and downregulated genes were defined as those with a normalized.avg_logFC (KO vs. WT) > 0.5 and p_adj_ < 0.05 or with a normalized.avg_logFC (KO vs. WT) < -0.5 and p_adj_ < 0.05, respectively.

**Fig. S7** Vascular networks comprised of WT and *CCM* KO cells show structural differences compared to WT controls. Vascular networks were prepared by mixing AICS-0054 WT (mTagRFPT) hiPSCs with AICS-0036 *CCM1*, *CCM2,* or *CCM3* KO hiPSCs in a 19:1 ratio and performing vascular network differentiation. **A** Staining for endothelial cells (CD31; blue) and pericytes (PDGFR-β; red) shows that mEGFP-tagged KO cells are part of both cell types. Furthermore, the staining shows that mosaic vascular networks containing *CCM3* KO cells are composed of much more convoluted endothelial and pericyte networks. **B** VE-cadherin staining of WT/KO mosaic networks demonstrates that most KO cells do not express VE-cadherin (scale bar = 50 µm).

**Video S1 – S4** 3D reconstructions of mosaic vascular networks show that most green-labeled *CCM1* (S1), *CCM2* (S2), and *CCM3* KO cells (S3) do not express VE-cadherin (red) compared to the green-labeled WT control cells (S4). Vascular networks were generated by mixing AICS-0054 WT (mTagRFPT) hiPSCs with AICS-0036 *CCM1*, *CCM2, CCM3* KO, or WT hiPSCs in a 19:1 ratio and performing vascular network differentiation. 3D reconstructions of the green-labeled AICS-0036 derived cells and VE-cadherin stainings (red) were created with FIJI v.1.54 (S1 = *CCM1* KO, S2 *= CCM2* KO, S3 = *CCM3* KO, S4 = WT control).

