## Supplementary figures and images for "High-throughput differentiation of human blood vessel organoids reveals overlapping and distinct functions of the cerebral cavernous malformation proteins"

### Supplementary Figure 1

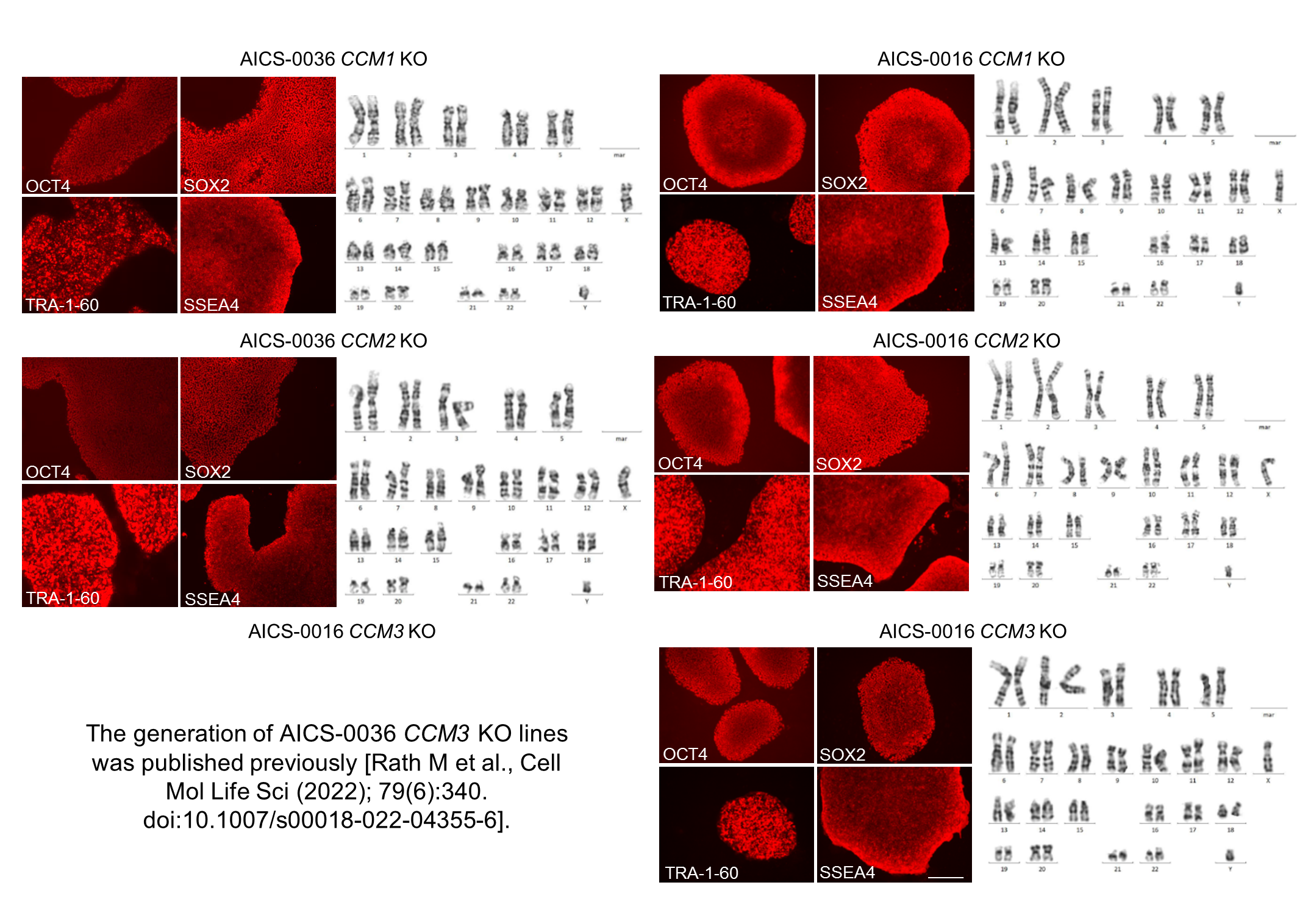

### Supplementary Figure 2

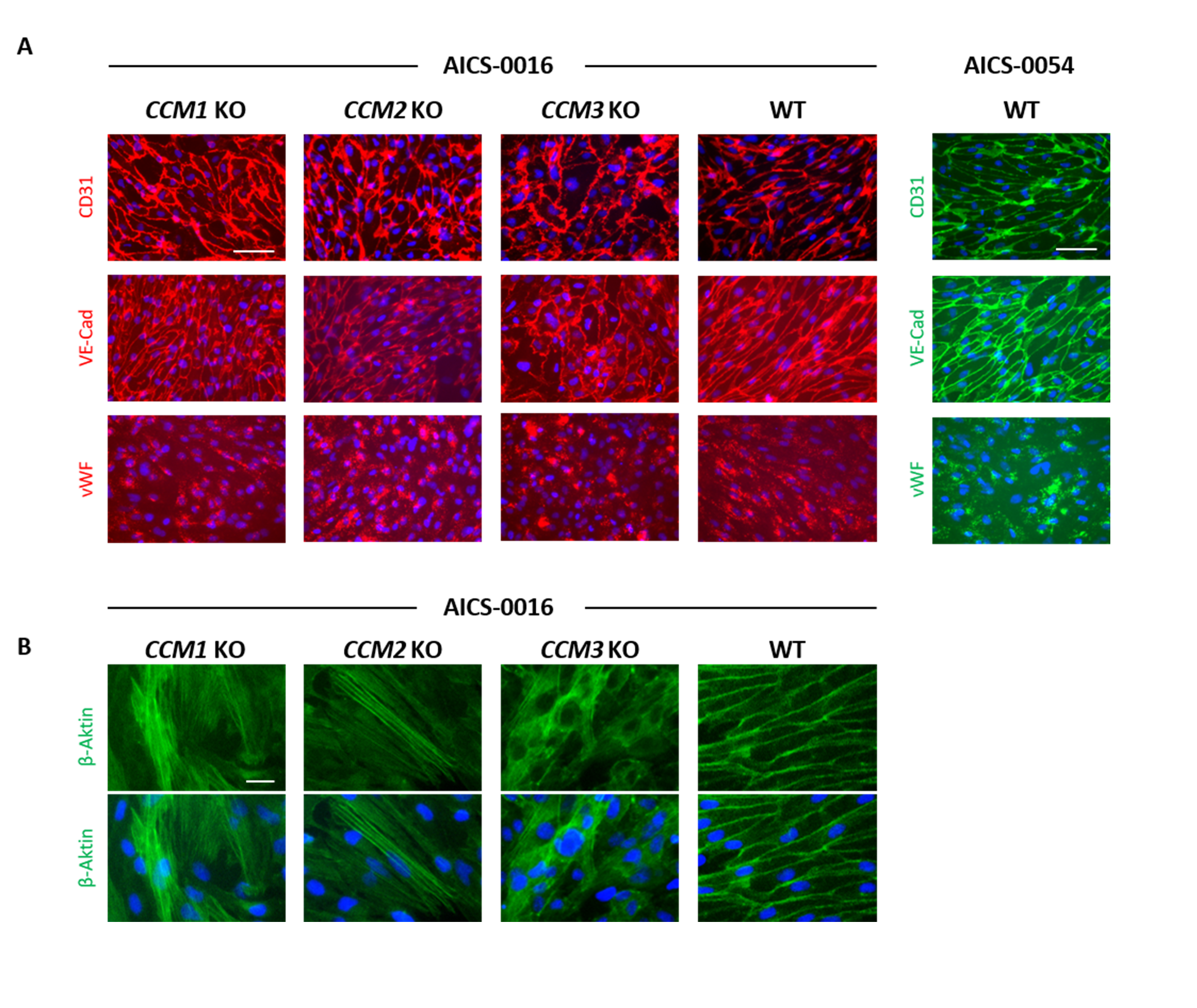

### Supplementary Figure 3

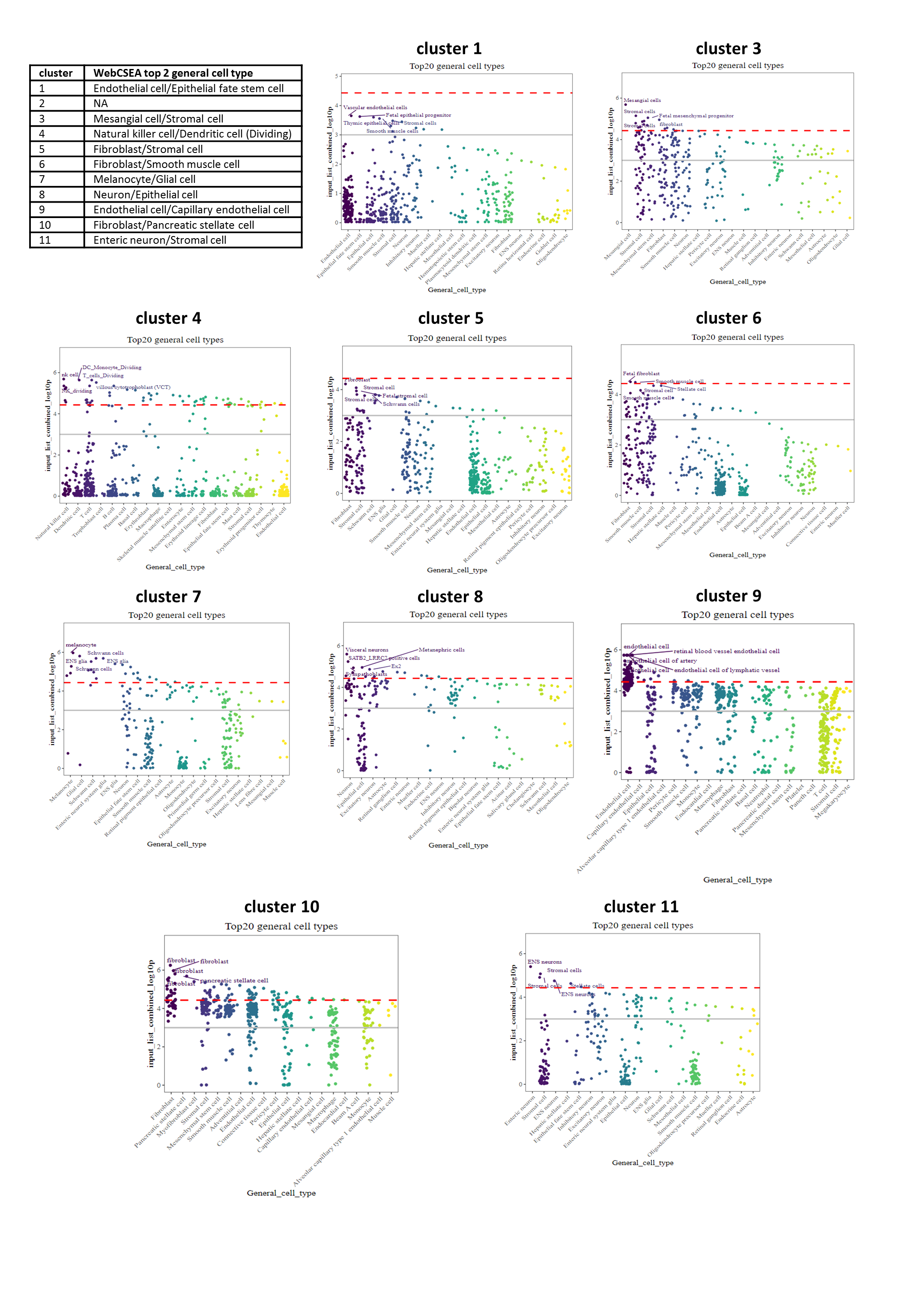

### Supplementary Figure 4

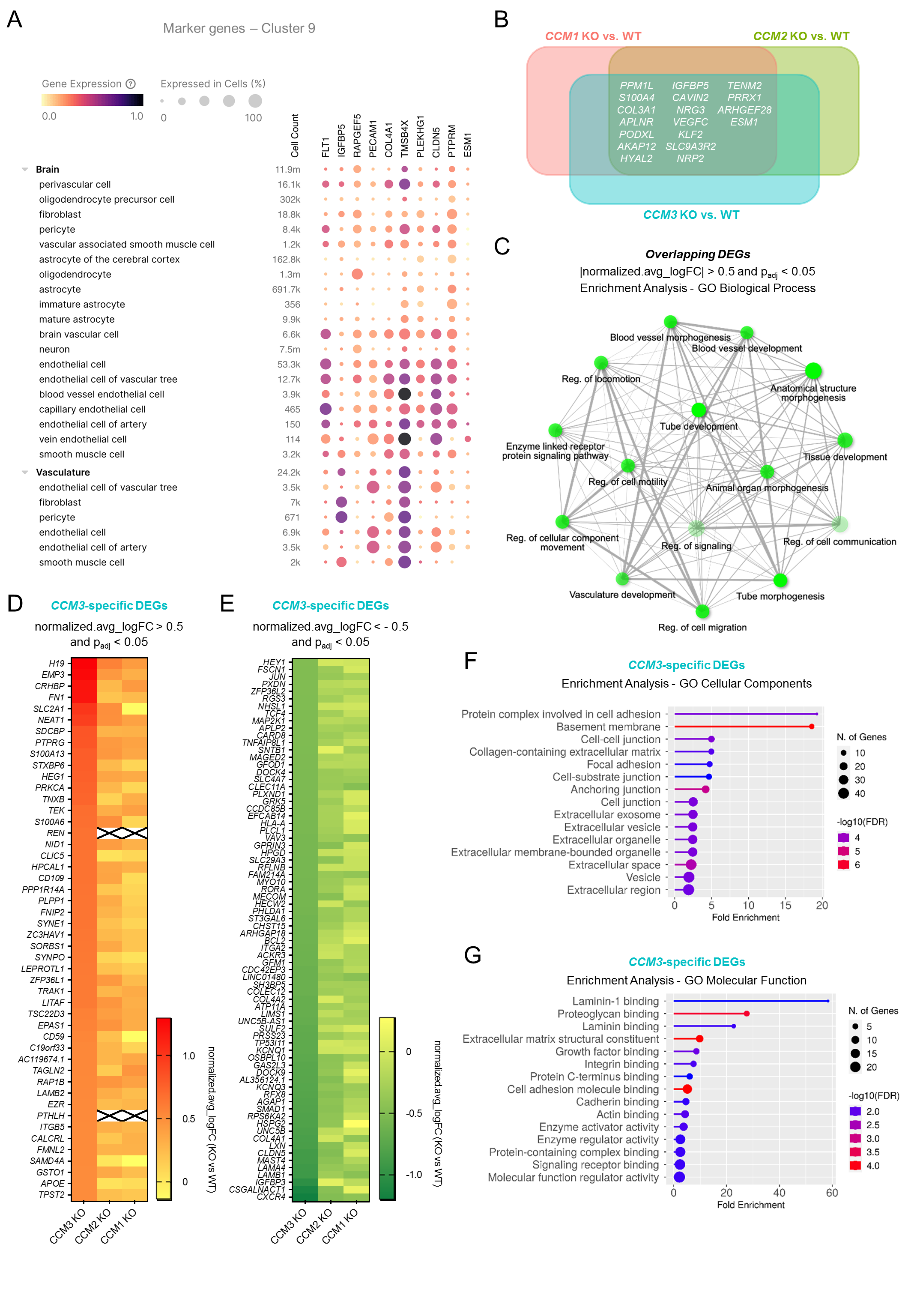

### Supplementary Figure 5

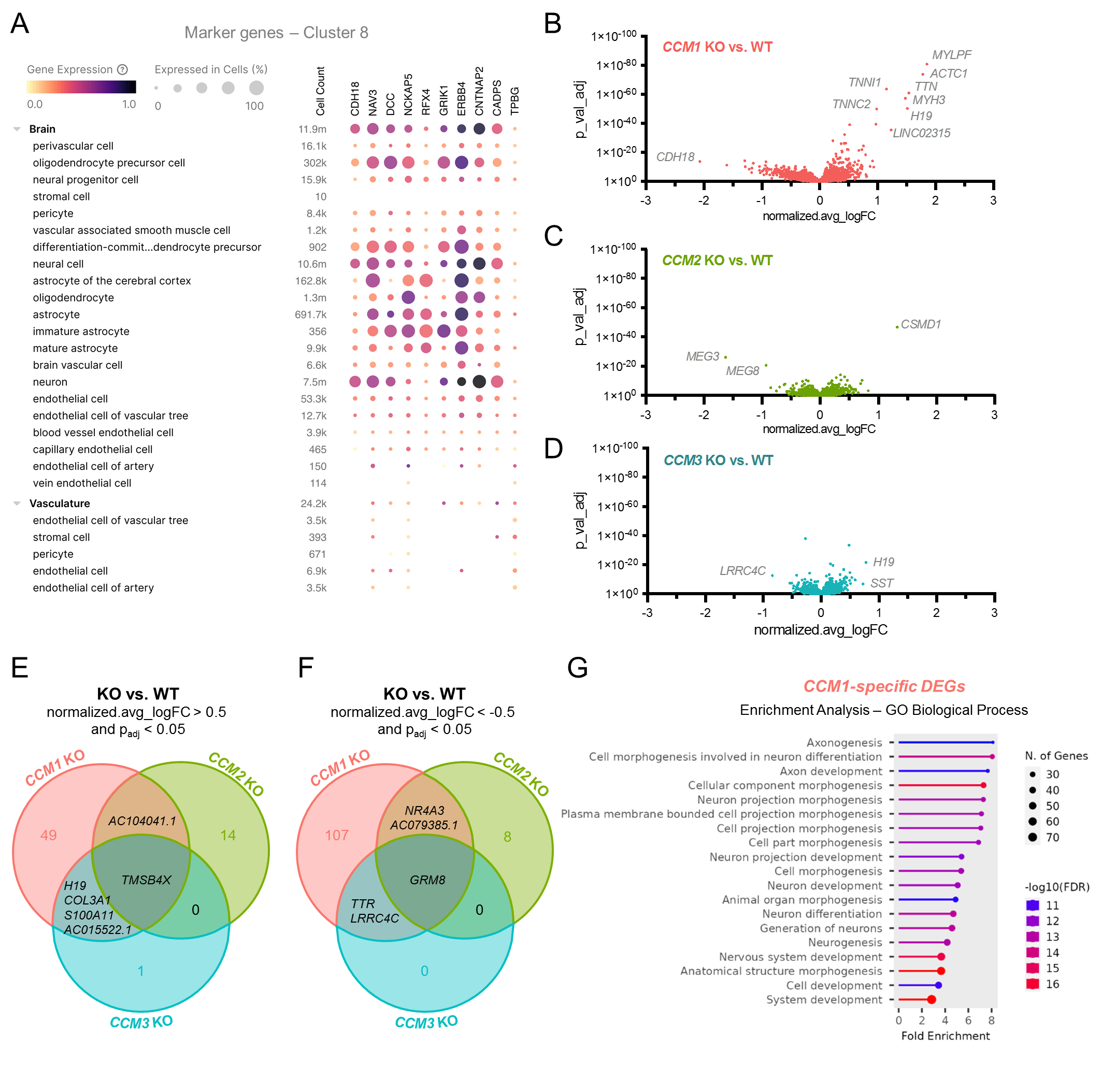

### Supplementary Figure 6

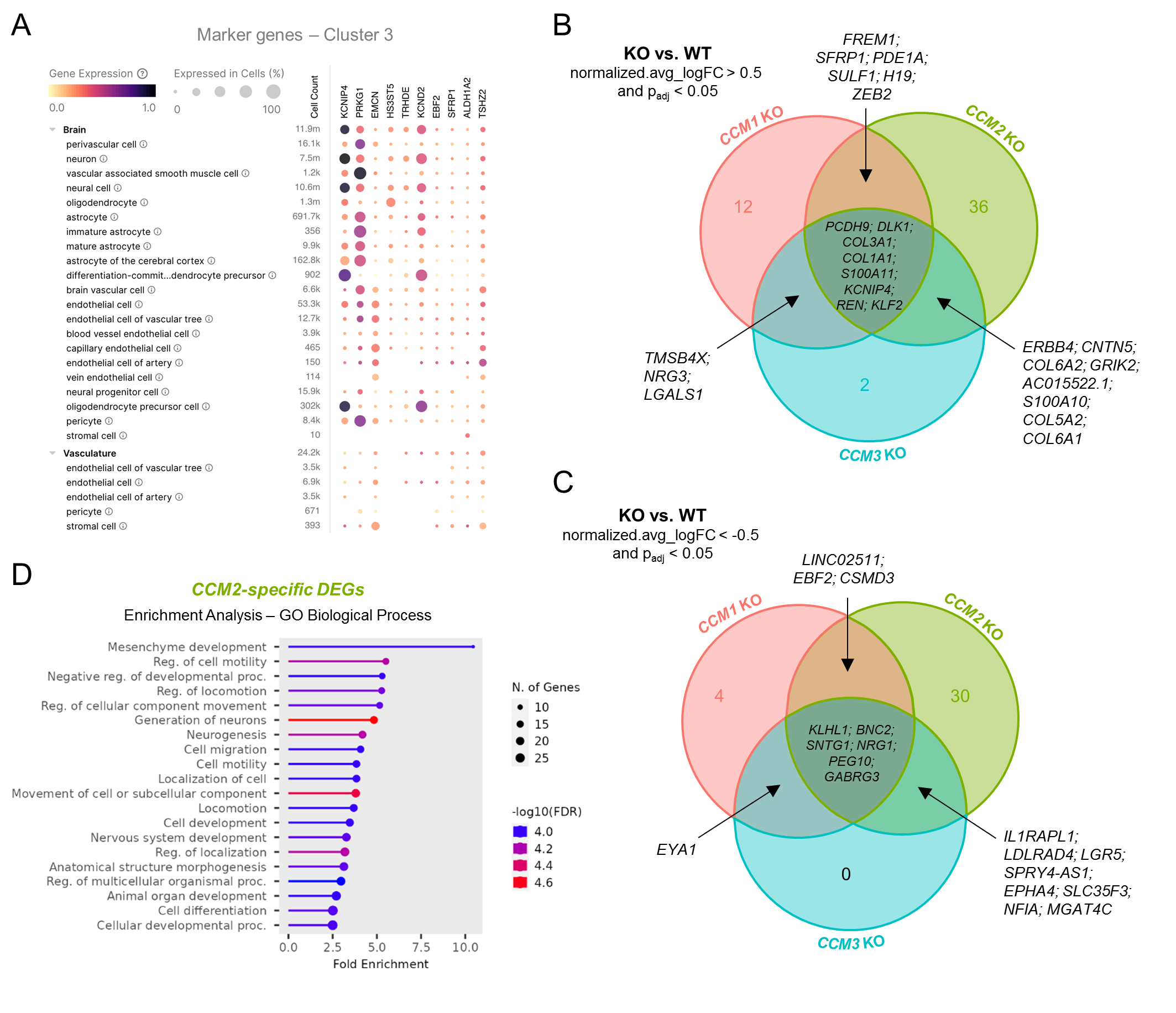

### Supplementary Figure 7

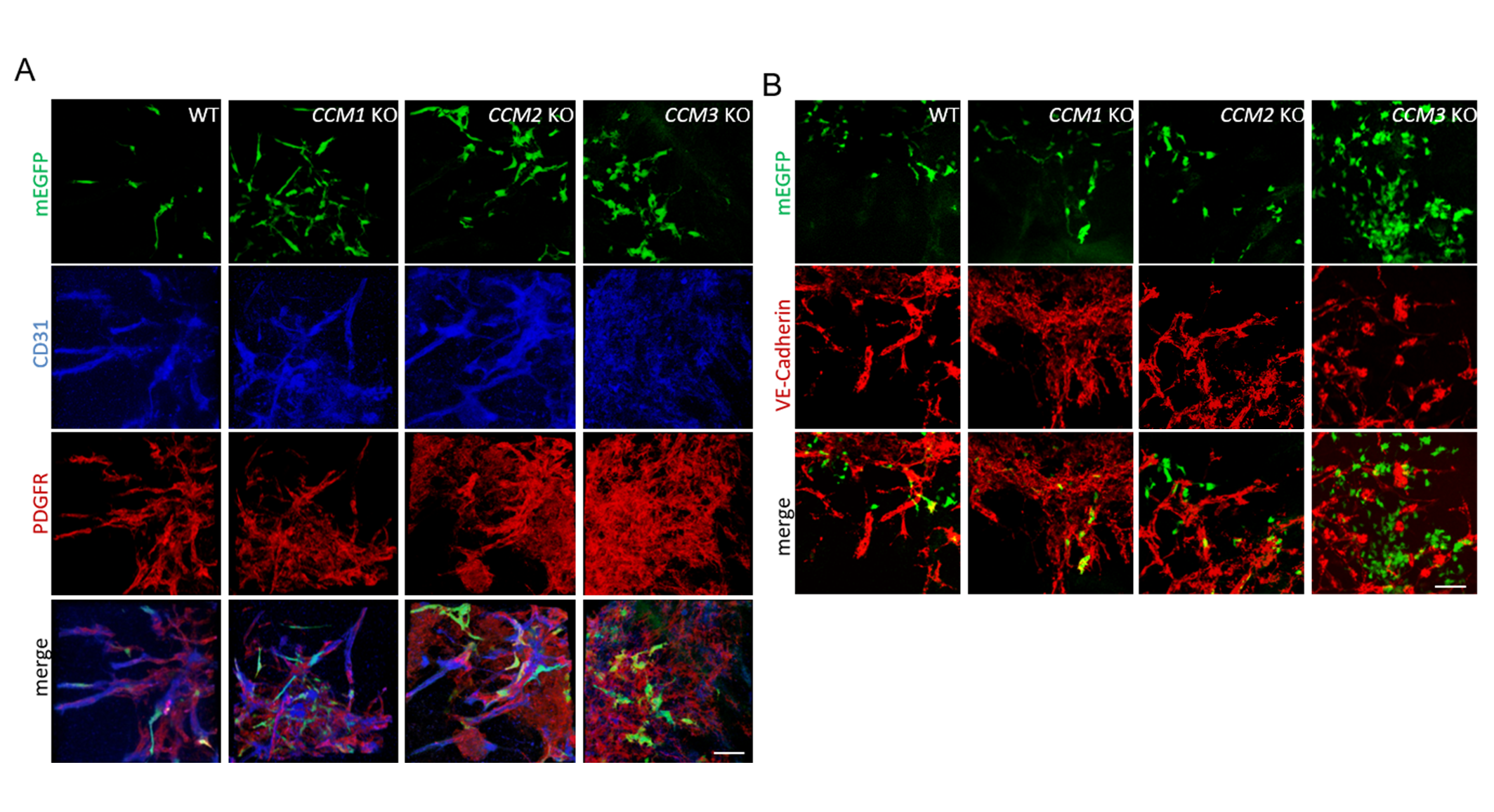

### Supplementary Video 1

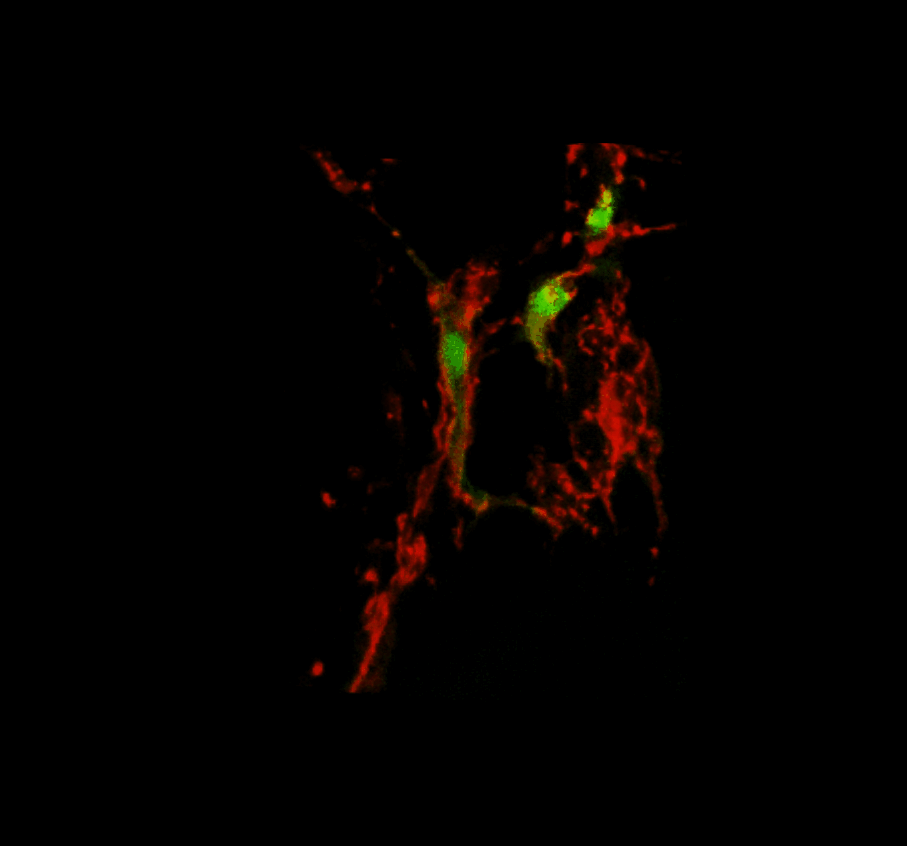

### Supplementary Video 2

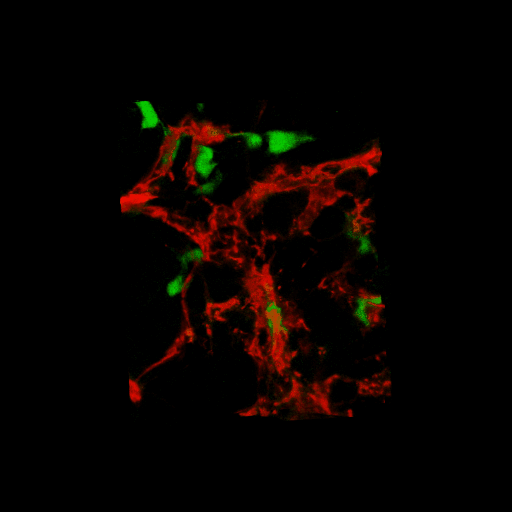

### Supplementary Video 3

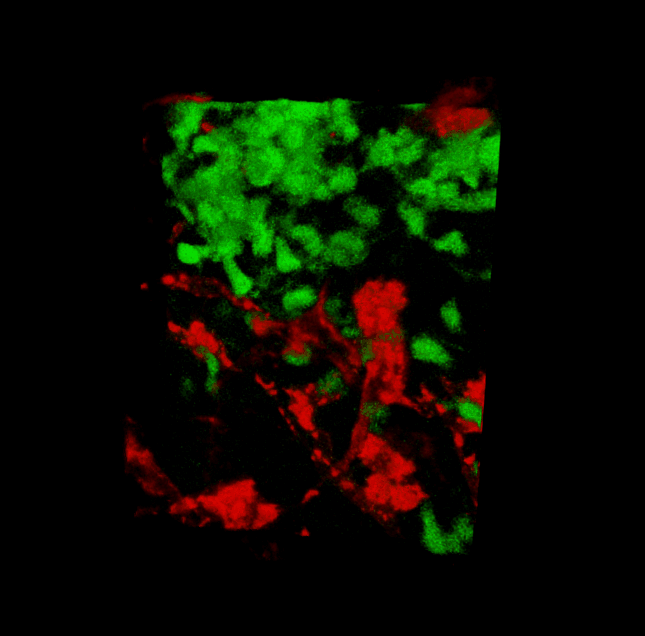

### Supplementary Video 4

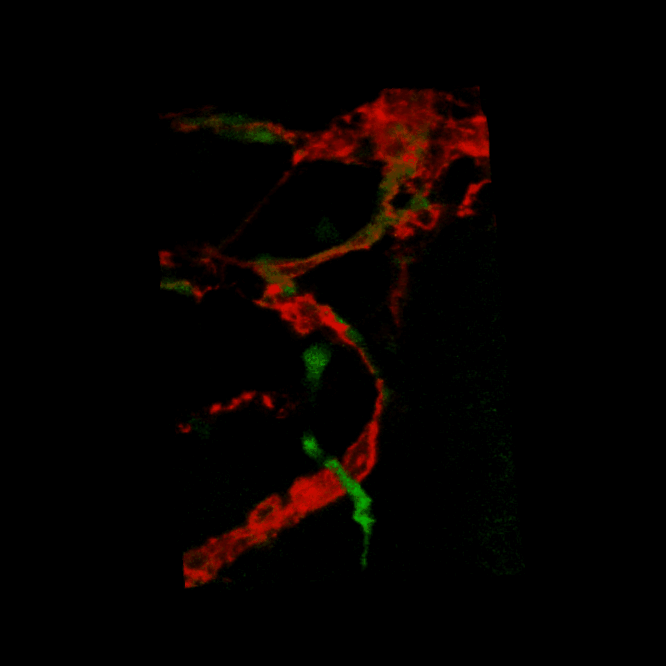
